# Ecological Inducers of the Yeast Filamentous Growth Pathway Reveal Environment-Dependent Roles for Pathway Components

**DOI:** 10.1101/2023.05.25.542306

**Authors:** Matthew D. Vandermeulen, Paul J. Cullen

## Abstract

Signaling modules, such as MAPK pathways, are evolutionarily conserved drivers of cell differentiation and stress responses. In many fungal species including pathogens, MAPK pathways control filamentous growth, where cells differentiate into an elongated cell type. The convenient model budding yeast *Saccharomyces cerevisiae* undergoes filamentous growth by the filamentous growth (fMAPK) pathway; however, the inducers of the pathway remain unclear, perhaps because pathway activity has been mainly studied in laboratory conditions. To address this knowledge gap, an ecological framework was employed, which uncovered new fMAPK pathway inducers, including pectin, a material found in plants, and the metabolic byproduct ethanol. We also show that induction by a known inducer of the pathway, the non-preferred carbon source galactose, required galactose metabolism and induced the pathway differently than glucose limitation or other non-preferred carbon sources. By exploring fMAPK pathway function in fruit, we found induction of the pathway led to visible digestion of fruit rind through a known target, *PGU1*, which encodes a pectolytic enzyme. Different stimuli revealed different modes of pathway signaling. For example, combinations of inducers (galactose and ethanol) stimulated the pathway to near maximal levels, which showed dispensability of several fMAPK pathway components (e.g. mucin sensor, PAK), but not others (e.g. adaptor, MAPKKK) and required the Ras2-PKA pathway. This included a difference between the transcription factor binding partners for the pathway, as Tec1p, but not Ste12p, was partly dispensable for fMAPK pathway activity. Thus, by exploring ecologically-relevant stimuli, new modes of MAPK pathway signaling were uncovered, perhaps revealing how a pathway can respond differently to specific environments.

**Data Availability Statement:** All data are in the manuscript and/or supporting information files.

## INTRODUCTION

Organisms can sense and respond to different signals in the environment. One of the ways this occurs is through the action of signal transduction pathways, such as evolutionarily conserved mitogen-activated protein kinase (MAPK) pathways [e.g. ERK-type, JNK-type, and p38-type (*1–3*)]. MAPK pathways sense and relay the signals from external/internal environments to induce a response, which typically occurs by the induction of gene expression (*4–8*). Much interest surrounding MAPK pathways comes from studies of their mis-regulation in diseases like cancer (*9–11*). With some exceptions, many aspects of MAPK pathways are still poorly defined like the fact that the stimuli that trigger pathways remain in many cases mysterious.

In addition to their role in animals, MAPK pathways also regulate biological responses in plants (*12, 13*) and fungi (*14–16*). In fungi, including single-celled yeasts, MAPK pathways can promote a cell differentiation response called filamentous growth. In pathogenic yeast, like the human pathogen *Candida albicans* (*17, 18*), and the plant pathogen *Ustilago maydis* (*19*), filamentous growth is controlled by ERK-type MAPK pathways and is a critical phenotype linked to virulence. During filamentous growth, cells grow in elongated tube-like structures (i.e. hyphae or pseudohyphae), and express specific cell adhesion molecules which promote attachment to surfaces and invasion into the host. In *U. maydis* [and other plant fungal pathogens (*20–23*)], MAPK pathways respond to cues from the plant surface to initiate invasion (*24, 25*). Although intensively studied, how fungal cells recognize, attach, and invade diverse environments through the action of signaling pathways remains an open question.

Certain aspects of filamentous growth and its regulation, including regulatory signaling pathways, are evolutionarily conserved between fungal species (*26, 27*). The model organism and saprotrophic budding yeast *Saccharomyces cerevisiae* undergoes filamentous growth (*28*) and has emerged as a useful model because of the ease of genetic manipulation and many tools available. In *S. cerevisiae* (*28, 29*) and *C. albicans* (*18, 30, 31*) filamentous growth occurs when cells encounter nutrient limitation. Quorum sensing molecules also influence filamentous growth and include alcohols like ethanol and butanol in *S. cerevisiae* (*32–35*) and farnesol and tyrosol in *C. albicans* (*36, 37*). *S. cerevisiae* do not form true hyphae but undergo a change in morphology from round yeast-form cells to adhesion-linked “chains” of elongated cells to produce filament-like structures that can invade into surfaces (i.e. invasive growth), presumably as a scavenging response [**Fig 1A**, (*28, 29, 38*)]. Also, *S. cerevisiae* (*39–41*), like *U. maydis* and many other plant-associated fungi (*42–44*), secrete a pectinase [Pgu1p in *S. cerevisiae*, an endo-polygalacturnase (*45*)] during filamentous growth to break down pectin (**Fig 1A**), which is a major component of plant cell walls (*46–49*).

**Figure 1.**
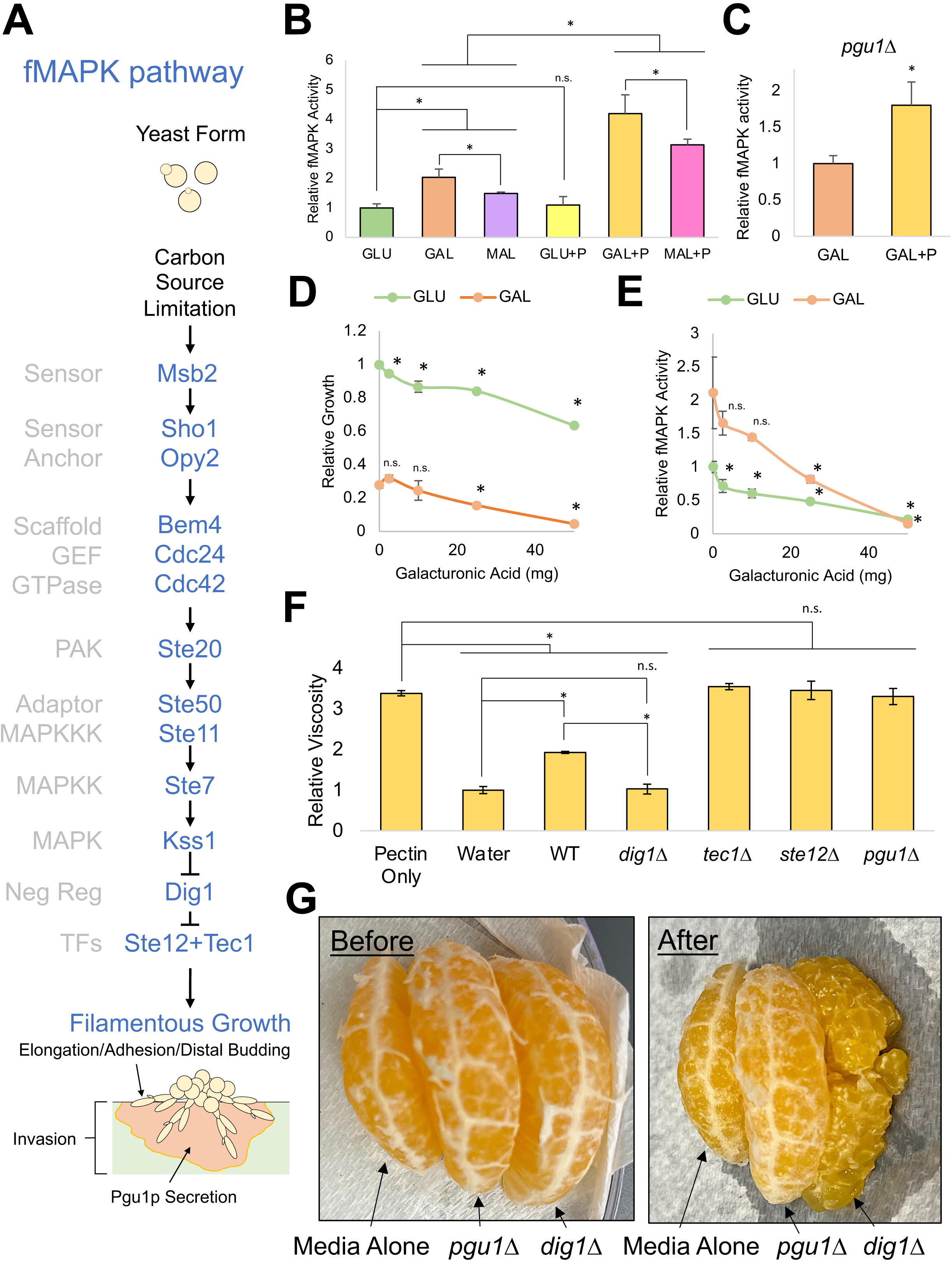
The fMAPK pathway is induced by pectin and regulates pectin breakdown in fruit. **A)** The fMAPK pathway regulates the switch to filamentous growth upon carbon source limitation. **B)** ß-galactosidase (lacZ) assays. Wild-type cells (PC313) were grown for 24 h in the in 5 mL of synthetic media with indicated carbon source. +P = + 1% pectin. Average relative fMAPK pathway activity for at least 3 replicates is reported, with values for GLU set to 1. Error bars represent standard deviation. Asterisk, p-value < 0.05 by Student’s t-test compared to the indicated condition. **C)** Same experiment as panel 1B except in the *pgu1Δ* mutant (PC7833). **D)** Relative growth of wild-type cells (PC313) in 5 mL synthetic media with the indicated carbon source after growth for 18 h. Galacturonic acid was added as indicated (mg). Average relative growth determined by OD_600_ across at least 3 replicates is reported, with values for GLU set to 1. Error bars represent standard deviation. Asterisk, p-value < 0.05 by Student’s t-test comparing galacturonic acid to carbon source alone. **E)** Same as panel 1D except ß-galactosidase (lacZ) assay for relative fMAPK pathway activity. **F)** Viscosity assay. Wild type cells (PC313) and the *pgu1Δ* (PC7833), *tec1Δ* (PC7675), *ste12Δ* (PC5651), and *dig1Δ* (PC7676) mutants were grown in GAL+P (1% pectin) solution for 17 h. Tube sedimentation was measured and compared to cultures of water and GAL+P (pectin only) with no cells added as controls for pectin breakdown and no pectin breakdown, respectively. Average relative viscosity from 3 replicates are reported, with values of tube sedimentation in water set to 1. Error bars represent standard deviation. Asterisk, p-value < 0.05 by Student’s t-test compared to indicated condition/strain. **G)** Pectin digestion. Mandarin orange wedges were incubated in supernatants of *pgu1Δ* and *dig1Δ* mutants for 24 h, which were derived from a 24h growth culture of cells in YPGAL, or YPGAL alone as a control. Left, wedges before incubation. Right, wedges after incubation.

Among several MAPK pathways in yeast (*50–52*), the filamentous growth MAPK (fMAPK) pathway is the main regulator of filamentous growth [(*28, 53–56*), **Fig 1A**]. The fMAPK pathway is headed by Msb2p, which is a member of the mucin family of signaling glycoproteins (*57–62*). Msb2p interacts with the tetraspan sensor protein, Sho1p (*61, 63–67*), and a single pass transmembrane protein, Opy2p, that anchors cytosolic proteins to the plasma membrane (*64, 68, 69*). Msb2p and Sho1p control the activity of the guanine nucleotide exchange factor (GEF) Cdc24p, which is the main activator (*70–72*) of the ubiquitous small Rho GTPase Cdc42p (*73–75*). In the context of filamentous growth, activation of the Cdc42p module requires at least three proteins, the bud-site GTPase Rsr1p (*76*), the polarity adaptor Bem1p (*77–79*) and the scaffold protein Bem4p (*77, 80*). Once activated, Cdc42p binds to and activates the p21-activated kinase (PAK), Ste20p (*67, 81–83*). Ste20p itself phosphorylates the MAPKKK, Ste11p [which is recruited to the plasma membrane by Opy2p and another adaptor Ste50p (*84, 85*)]. At the plasma membrane, Ste11p phosphorylates the MAPKK, Ste7p, which then phosphorylates the MAPK, Kss1p (*81, 86–89*). The MAP kinase Kss1p regulates two transcription factors, Ste12p and the TEA/ATTS-type Tec1p (*86, 90–93*), which dimerize to co-regulate the expression of target genes that bring about filamentous growth. Ste12p and Tec1p also work with other transcriptional regulators, including the co-activators Msa1p and Msa2p (*92*), and the strong transcriptional repressor, Dig1p (*87, 88, 94*).

Unlike pathogens, where ecology in the host takes center stage, studies of the fMAPK pathway in *S. cerevisiae* have mainly been performed in laboratory conditions. However, *Saccharomyces* yeast are commonly found in wild and domesticated habitats, including fruits, tree sap and bark, insect vectors, leaf litter, soil, and rotten wood (*95–105*). Signaling pathways in *S. cerevisiae* are likely to have evolved to sense and respond to stimuli from these diverse environments. To explore this possibility, we investigated fMAPK pathway activation by compounds that might be expected to be encountered in the wild. We considered plant-associated compounds, like pectin, and the metabolic byproduct and quorum-sensing molecule, ethanol, which is also an inhibitor of microbial competitors (*106, 107*), and an attractant for insect vectors (*102, 108–111*). We also examined carbon sources like galactose more closely, which is abundant in natural habitats of *S. cerevisiae* (*101, 102, 112*), including forest leaf litter/soil (*104, 113*), and certain fruits, like persimmons (*114–116*).

This ‘ecological approach’ uncovered new inducers of the fMAPK pathway, including pectin and ethanol. We also found that galactose induced fMAPK pathway activity differently than glucose limitation or other non-preferred carbon sources. Combinations of commonly encountered inducers (galactose with ethanol) activated the pathway to near maximal levels, which has not been previously observed in laboratory settings. Maximal activation of the pathway partly by-passed the requirement of several core components of the pathway (Msb2p, Sho1p, Opy2p, Ste20p, Bem4p and Tec1p) but not others (Ste50p, Ste11p, and Ste12p). We also identified a critical role for the Ras2-PKA pathway in fMAPK pathway regulation in response to ethanol. Thus, studying a model organism from an ecological perspective can provide insights into pathway regulation that may generally apply to other systems, like pathogens who thrive in the unique ecologies of their hosts.

## RESULTS

### Pectin is a new inducer of the fMAPK pathway

The fMAPK pathway regulates pectinase levels (*41*); therefore, we tested whether the fMAPK pathway is activated when cells encounter plant material/compounds to promote pectin breakdown. Several plant compounds were tested, including pectin, breakdown products of pectin (di-galacturonic acid and galacturonic acid), and indoleacetic acid (IAA), a plant hormone previously shown to stimulate filamentous growth (*117*). The activity of the fMAPK pathway was measured in a filamentous strain (∑1278b background) by a transcriptional reporter [p*FRE-lacZ*, (*118*)]. To test the effect of pectin on fMAPK pathway activity, a 1% pectin solution was made in media containing the preferred carbon source glucose, or non-preferred carbon sources, galactose or maltose. In glucose, the activity of the fMAPK pathway was not stimulated by pectin (**Fig 1B**, compare GLU to GLU+P). However, pectin stimulated the fMAPK pathway in media supplemented with galactose or maltose (**Fig 1B**, compare GAL to GAL+P and MAL to MAL+P). The fact that pectin induced the fMAPK pathway only in the absence of glucose may suggest that pectin induction is subject to glucose repression (*119*).

Pectin is a polymer composed of chains of galacturonic acid and other sugars (*48*). Pectin, and its breakdown product by the Pgu1p enzyme, di-galacturonic acid, both regulate *PGU1* expression (*120, 121*); therefore, pectin may be recognized by the fMAPK pathway as a polymer or by its breakdown products. Unlike pectin, di-galacturonic acid did not induce the fMAPK pathway (*Fig S1A*). In cells lacking pectinase (*pgu1Δ*), where pectin breakdown does not occur (see below), pectin induced the fMAPK pathway (**Fig 1C**, *pgu1Δ* mutant shows similar increase between GAL and GAL+P as WT in **Fig 1B**). Thus, the fMAPK pathway is induced by pectin and does not require breakdown products of Pgu1p activity.

Presumably, *S. cerevisiae* does not break down di-galacturonic acid into galacturonic acid because galacturonic acid inhibits growth (*122*). Some fungal and bacterial organisms other than *S. cerevisiae* can break down pectin to galacturonic acid to use as a carbon source (*123–126*). This body of data suggests that galacturonic acid may be encountered in the wild. As expected, galacturonic acid inhibited the growth of yeast cells (**Fig 1D**), however, it also caused a reduction in fMAPK pathway activity (**Fig 1E**). The reduced fMAPK pathway activity is likely not due to growth inhibition, as growth inhibition by other compounds did not cause a reduction in fMAPK pathway activity (see below, ethanol). The plant hormone, IAA, did not affect fMAPK pathway activity (*Fig S1B*). Collectively, these experiments identify galacturonic acid as an inhibitor and pectin as a new inducer of the fMAPK pathway.

### A function for the fMAPK pathway in pectin degradation in fruit

Yeast cells may break down pectin to improve accessibility to plant tissues, and therefore nutrients, as has been previously suggested (*39–41, 127*). Pectin may also be broken down to use as a carbon source. This latter possibility seems unlikely because *S. cerevisiae* was not able to grow in pectin as the sole carbon source (WT OD_600_ in synthetic medium with 2% pectin as carbon source remained < 0.08 after 16 h). Therefore, we focused on testing whether pectin breakdown might improve accessibility to the plant environment.

Pectin breakdown by Pgu1p has previously been visualized by a plate-based test that measures enzymatic activity (*41, 128*). Here, to more directly test whether pectin breakdown affects accessibility to the plant environment, two tests were developed. The first test was based on the fact that pectin-rich solutions are viscous, which reflects pectin acting as a physical barrier to yeast cells; therefore, we measured changes in viscosity of pectin solutions after incubation with wild-type cells and fMAPK pathway mutants. Viscosity can be measured by determining the time for a weight to reach the bottom of a solution. The viscosity of a 1% pectin solution was measured as a control (**Fig 1F**, pectin only) and compared to water, a control for complete pectin breakdown (**Fig 1F**, water). The viscosity of a pectin solution was found to be reduced after a 17 h incubation with wild-type cells (**Fig 1F**, WT). The reduction in viscosity was dependent on pectinase activity, as seen in cells lacking Pgu1p (**Fig 1F**, *pgu1*∆). The reduction in viscosity was also dependent upon the fMAPK pathway as seen in cells lacking transcription factors Tec1p or Ste12p (**Fig 1F**, *ste12*∆ or *tec1*∆). Viscosity was strongly reduced when incubated with cells lacking the negative regulator Dig1p, which has elevated fMAPK pathway activity (**Fig 1F**, *dig1Δ*). Therefore, the fMAPK pathway functions to reduce viscosity of pectin-based solutions, and may facilitate accessibility to plant-based environments.

In the second test, pectin levels were examined in rinds on a fruit (mandarin oranges), an environment where *S. cerevisiae* has been isolated (*101, 112*). Mandarin orange wedges were incubated for 24 h in supernatants derived from the *dig1Δ* and *pgu1Δ* mutants and compared to incubation with medium alone (**Fig 1G**, more images in *Fig S2*). This result showed that the fMAPK pathway and Pgu1p cause a detectable loss of pectin material in fruit rind. Moreover, it was clear that the fruit tissue was damaged by the *dig1Δ* mutant (**Fig 1G**, after), which one would expect to promote the liberation of nutrients. Thus, one function of the fMAPK pathway is to deteriorate pectin in the rind that connects plant tissues to liberate nutrients from fruits.

### Galactose induces the fMAPK pathway in a separate way than glucose limitation or other carbon sources

Growth in the non-preferred carbon source galactose stimulates the fMAPK pathway (*129, 130*), which may be due to the absence of a preferred carbon source like glucose. Alternatively, galactose may specifically induce the fMAPK pathway. Due to the prevalence of galactose in yeast habitats, we tested galactose in comparison to other sugars for induction of the fMAPK pathway.

Cells were examined for growth, fMAPK pathway activity, and cell morphology in preferred carbon sources where cells grew well [**Fig 2A**, Glucose (GLU) and Fructose (FRU)], and non-preferred carbon sources where cells grew similarly and poorly [galactose (GAL), sucrose (SUC), maltose (MAL), and glycerol (GLY)]. The fMAPK pathway was induced by galactose to higher levels than by other carbon sources (**Fig 2B**, 8 h). Intriguingly, cells became elongated in galactose, sucrose, and maltose but not glucose, fructose, or glycerol (**Fig 2C**), indicating that the morphological changes associated with filamentous growth, and the activity of the fMAPK pathway can occur separately. Glycerol failed to induce filamentous growth or the fMAPK pathway, indicating that some poor carbon sources do not trigger this differentiation response, at least at the time point tested here. At 24 h, cells grown in maltose and galactose grew similarly (*Fig S3A*), and galactose and maltose both induced the fMAPK pathway relative to glucose, although galactose induced the pathway to higher levels (**Fig 2B**). Therefore, galactose induced the fMAPK pathway at an earlier time point relative to other carbon sources (8 h, **Fig 2B**) and to higher levels at a later timepoint (24 h, **Fig 1B**). Thus, induction of the fMAPK pathway occurs by the addition of galactose in a manner that is distinct from other poor carbon sources.

**Figure 2.**
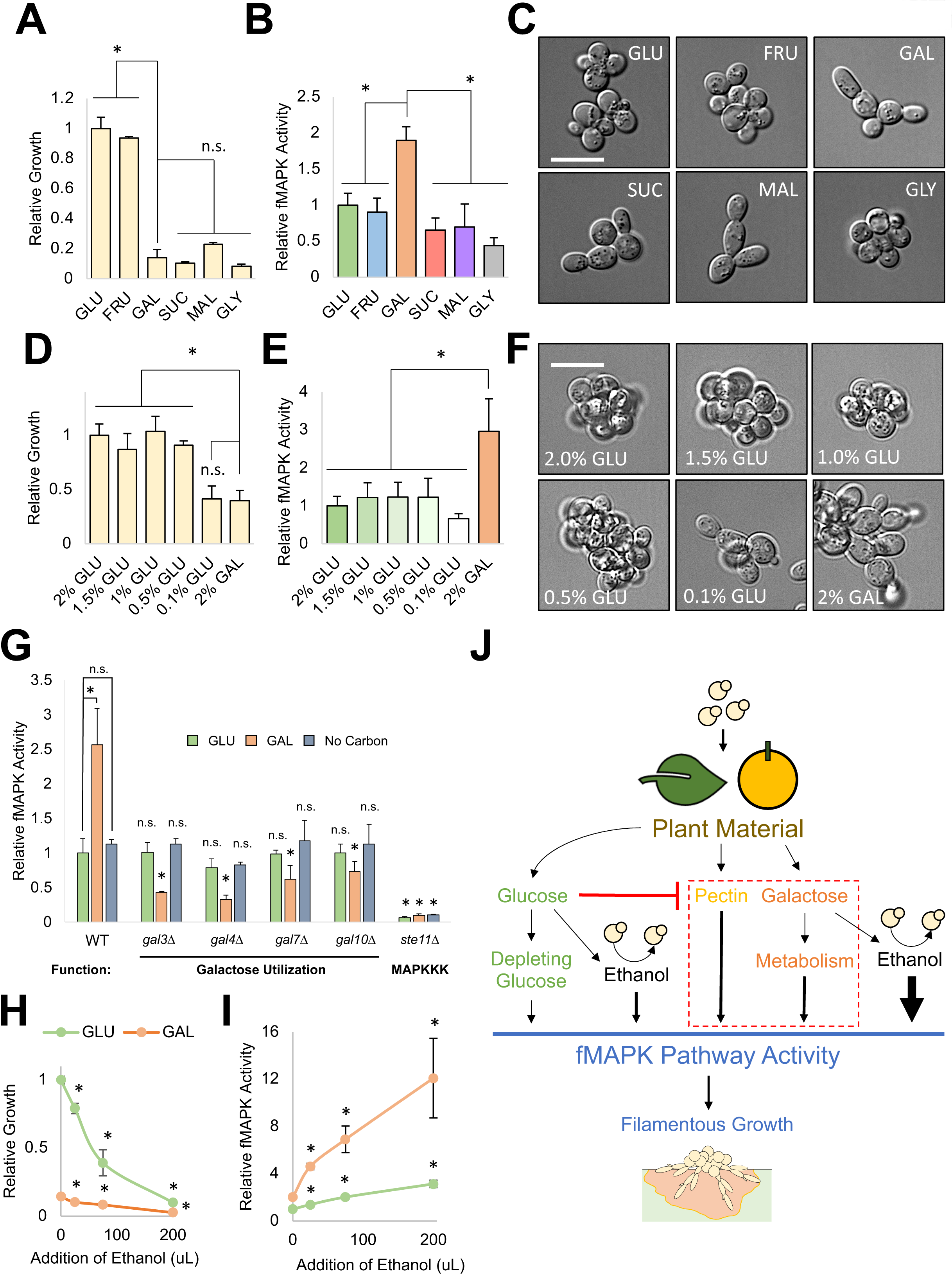
Galactose and ethanol induce the fMAPK pathway. **A-C)** Wild-type cells (PC313) were grown in 5 mL synthetic media with indicated carbon source for 8 h. <u>Panel A</u>, average relative growth determined by OD_600_ of at least 3 replicates are reported with GLU values set to 1. Error bars represent standard deviation. Asterisk, p-value < 0.05 by Student’s t-test compared to GAL. <u>Panel B</u>, ß-galactosidase (lacZ) assays, average relative fMAPK pathway activity across at least 3 replicates is reported, with GLU values set to 1. Error bars represent standard deviation. Asterisk, p-value < 0.05 by Student’s t-test to GAL. <u>Panel C</u>, images of cell morphology in indicated media. Bar = 10 um. **D-F)** Wild-type cells (PC313) were grown in 2 mL synthetic media with glucose or galactose at indicated concentrations for 15 h. <u>Panel D</u>, average relative growth determined by OD_600_ of at least 3 replicates are reported with 2% GLU values set to 1. Error bars represent standard deviation. Asterisk, p-value < 0.05 by Student’s t-test compared to 2% GAL. <u>Panel E</u>, ß-galactosidase (lacZ) assays, average relative fMAPK pathway activity across at least 3 replicates is reported, with 2% GLU values set to 1. Error bars represent standard deviation. Asterisk, p-value < 0.05 by Student’s t-test to 2% GAL. <u>Panel F</u>, images of cell morphology in indicated media. Bar = 10 um. **G)** ß-galactosidase (lacZ) assays. Wild-type (PC313) cells and the *ste11Δ* (PC5024), *gal3Δ* (PC7849), *gal4Δ* (PC7845), *gal7Δ* (PC7844), and *gal10Δ* (PC7846) were grown in 2 mL rich media (Yeast extract, peptone) with indicated carbon source (or no carbon source as a control) for 7 h. Average relative fMAPK pathway activity across at least 3 replicates is reported, with wild-type values in GLU set to 1. Error bars represent standard deviation. Asterisk for wild-type values, p-value < 0.05 by Student’s t-test compared to indicated condition. Asterisk for mutants’ values, p-value < 0.05 by Student’s t-test compared to wild-type values from same condition. **H-I)** Wild-type cells (PC313) were grown in glucose or galactose with the indicated concentrations of ethanol added to 5 mL cultures for 17 h. Max addition, 200ul of ethanol, is 3.85% ethanol. <u>Panel G</u>, average relative growth determined by OD_600_ of at least 3 replicates are reported with GLU values set to 1. Error bars represent standard deviation. Asterisk, p-value < 0.05 by Student’s t-test comparing tested ethanol concentration to its respective carbon source with no ethanol added. <u>Panel H</u>, ß-galactosidase (lacZ) assays, average relative fMAPK pathway activity across at least 3 replicates is reported, with values in GLU set to 1. Error bars represent standard deviation. Asterisk, p-value < 0.05 by Student’s t-test comparing tested ethanol concentration to the same carbon source with no ethanol added. **J)** Model of new inducers. Depleting glucose activates the pathway to a minor degree (smallest arrow). Pectin and galactose induce the fMAPK pathway in environments depleted for glucose. Ethanol induces the pathway in glucose and more strongly in galactose (galactose with ethanol has largest arrow). fMAPK pathway induction by these stimuli leads to filamentous growth.

We previously showed that glucose depletion triggers invasive growth (*29*). In line with this observation, fMAPK pathway activity increased over time in media containing glucose, presumably as glucose became depleted (*Fig S3B*). However, in addition to glucose depletion, the above findings suggest that the addition of galactose might induce the fMAPK pathway. To directly compare ‘glucose limitation’ to ‘galactose induction’, different concentrations of glucose were compared for growth (**Fig 2D**), fMAPK pathway activity (**Fig 2E**), and cell elongation (**Fig 2F**). Cells grew similarly and poorly in both 0.1% glucose and 2% galactose and showed similar cell elongation (**Fig 2, D** and **F**); however, galactose induced fMAPK pathway activity compared to glucose depletion (**Fig 2E**). Surprisingly, lower glucose concentrations did not cause an increase fMAPK pathway activity by this method (**Fig 2E**, compare 2% GLU to 0.1% GLU). This may be due to the fact that when cells deplete glucose over time there is a build-up of other signals in the environment (e.g. metabolites) that do not occur when cells are transferred to limiting glucose. Likewise, by examining fMAPK pathway activity at a time point when growth in galactose reached a comparable cell density as growth in glucose (*Fig S3A*, 24 h), and nutrients have presumably depleted naturally, galactose-dependent induction of the fMAPK pathway was higher than seen in glucose (**Fig 1B**). Thus, induction of the fMAPK pathway by galactose is different and more robust than induction of the pathway by glucose limitation.

A metabolic pathway is responsible for galactose uptake and utilization (*131*) and contains proteins for the transport of galactose into the cell [Gal2p, galactose permease, (*132, 133*)] and transduction of the galactose signal [through Gal3p and Gal80p, (*134, 135*)] to increase the activity of the transcription factor Gal4p, (*136, 137*). Gal4p induces the expression of genes encoding enzymes required for galactose metabolism [e.g. Gal7p and Gal10p, (*138–140*)]. Several galactose pathway mutants (*gal3Δ*, *gal4Δ, gal7Δ,* and *gal10Δ*) were constructed and tested for fMAPK pathway activation by galactose in reference to a mutant lacking a core component of the fMAPK pathway (*ste11Δ*). *GAL* genes were required for galactose-dependent induction of the fMAPK pathway (**Fig 2G**, orange). As expected, the *GAL* genes were not required for fMAPK pathway activity in medium with glucose as the carbon source or no carbon source (**Fig 2G**, green and blue). These results show that galactose metabolism is required for the induction of the fMAPK pathway by galactose.

### Ethanol induces the fMAPK pathway

Ethanol is a byproduct of glycolysis and an inducer of invasive growth (*32–34*). Like galacturonic acid (**Fig 1D**), ethanol induced a growth defect (**Fig 2H**, green). However, unlike galacturonic acid, ethanol stimulated fMAPK pathway activity (**Fig 2I**, green). Ethanol-dependent induction of the fMAPK pathway, like pectin, was more evident in media containing galactose than glucose (**Fig 2I**); however, unlike pectin, ethanol was able to induce fMAPK pathway activity in glucose to some degree (∼3-fold). In fact, induction of the fMAPK pathway by ethanol in glucose occurred to comparable levels as induction by galactose (**Fig 2I**, compare GLU with ethanol to GAL without ethanol). Thus, ethanol induces the fMAPK pathway. Collectively, the results presented here show that the fMAPK pathway can be induced by stimuli (pectin, galactose, and ethanol) that might be commonly encountered when cells are exposed to and metabolize plant material in natural settings. Taking into account the dependency on glucose limitation for several of the inducers, we suggest a model for how these stimuli might be encountered (**Fig 2J**).

### Combinations of inducers stimulate the fMAPK pathway to near maximal levels and reveal environment-dependent roles for pathway components

Multiple inducers might be encountered in the wild and were therefore tested in combinations. As shown above in galactose, pectin or ethanol stimulated fMAPK pathway activity above galactose alone (**Fig 3A**, WT). These results suggest that pectin and ethanol have additive effects with galactose. When galactose, pectin, and ethanol were combined there was no further increase in fMAPK pathway activity above galactose with ethanol (*Fig S4*), suggesting pectin does not necessarily have an additive effect with ethanol. Alternatively, the addition of pectin to galactose with ethanol may not further stimulate pathway activity because the pathway may be maximally activated. To test this hypothesis, wild-type cells were compared to the *dig1Δ* mutant, which shows very high fMAPK pathway activity seen in laboratory settings (*141*). The *dig1Δ* mutant showed elevated fMAPK pathway activity compared to wild-type cells in most conditions, except in galactose with ethanol (**Fig 3A**, compare wild type to the *dig1Δ* mutant, galactose with ethanol, striped orange) or when all three inducers were combined (*Fig S4*). These results show that the fMAPK pathway can be activated to near maximal levels when cells encounter a combination of inducers (**Fig 2J**, galactose with ethanol, largest inducing arrow).

**Figure 3.**
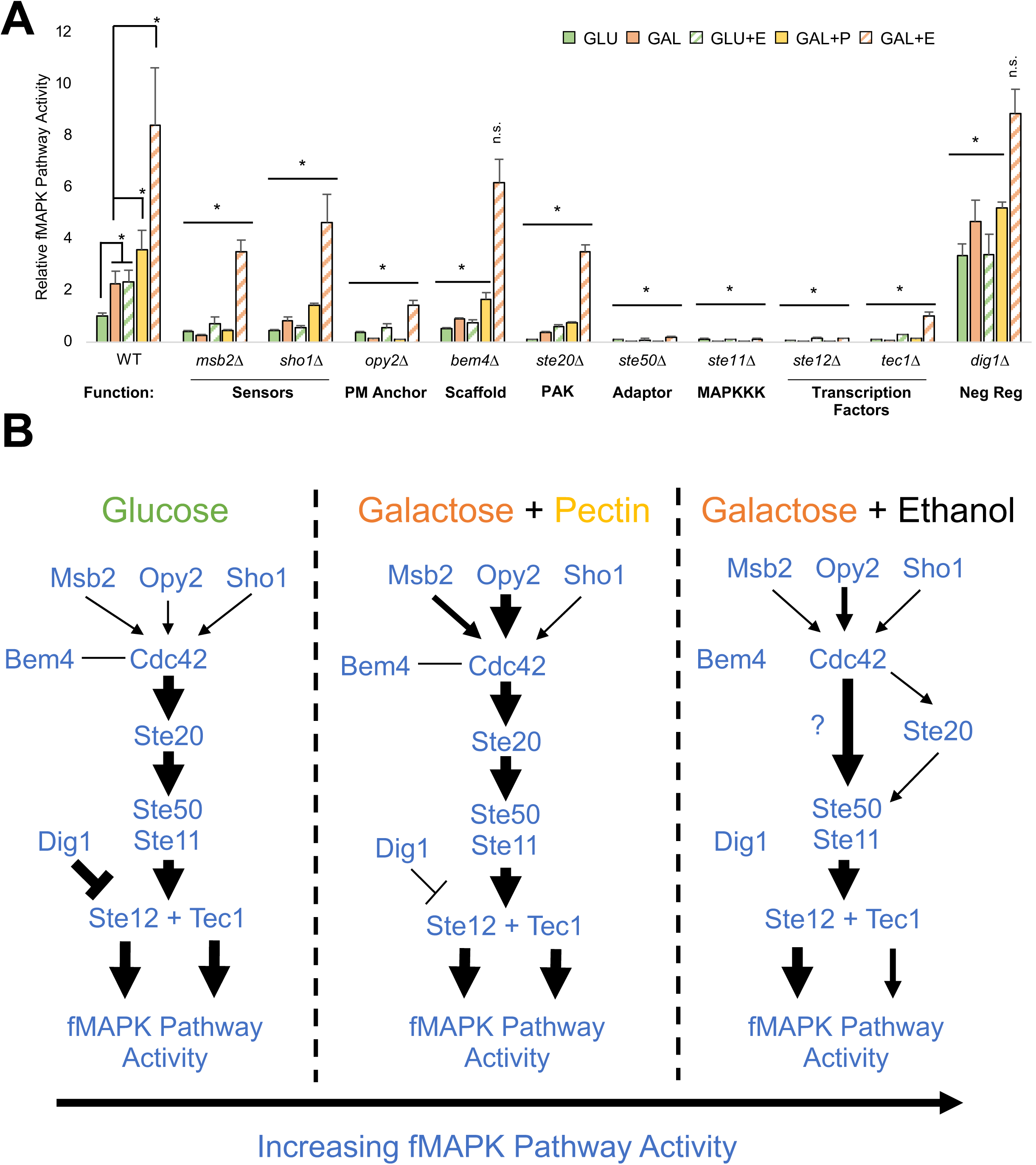
Combinations of inducers activates the fMAPK pathway to near maximal levels and reveals roles of fMAPK pathway components under different conditions. **A)** ß-galactosidase (*lacZ*) assays. Wild-type cells (PC313) and the *msb2Δ* (PC961), *opy2Δ* (PC3894), *sho1Δ* (PC5026), *bem4Δ* (PC3343), *ste20Δ* (PC7871), *ste50Δ* (PC4982), *ste11Δ* (PC5024), *ste12Δ* (PC5651), *tec1Δ* (PC7675), and *dig1Δ* (PC7676) mutants were grown in 2 mL synthetic media with indicated carbon source for 17 h. +P = + 1% pectin. +E = +3.85% ethanol. Average relative fMAPK pathway activity across at least 3 replicates is reported, with wild-type values in GLU set to 1. Error bars represent standard deviation. Asterisk for wild-type values, p-value < 0.05 by Student’s t-test compared to indicated condition. Asterisk for mutants’ values, p-value < 0.05 by Student’s t-test compared to wild-type values from same condition. **B)** Model depicting findings in panel A. Arrow size represents MAPK activity based on the data in panel 3A.

We next tested how different inducers may be sensed and relayed by fMAPK pathway components (**Fig 1A**). In general, cells lacking fMAPK pathway regulatory proteins (*msb2Δ, opy2Δ, sho1Δ, bem4Δ, ste20Δ, ste50Δ, ste11Δ, ste12Δ,* and *tec1Δ* mutants; *cdc24Δ* and *cdc42Δ* mutants are inviable and were not tested) showed reduced fMAPK pathway activity compared to wild type in response to the inducers tested (**Fig 3A**). However, several findings were worth noting. First, the proteins that reside at the plasma membrane, Msb2p, Sho1p, and Opy2p were more critical for signaling in some environments than others. For example, the *msb2Δ* mutant showed a stronger reduction in fMAPK pathway activity in galactose compared to galactose with ethanol relative to wild-type levels [**Fig 3A**, galactose, *msb2Δ* shows ∼10-fold-reduction compared to WT, galactose with ethanol, *msb2Δ* shows ∼2.5-fold-reduction compared to WT].

Second, Msb2p, Sho1p, and Opy2p showed different requirements for signaling compared to each other depending on the environment. For example, the *msb2Δ*, *sho1Δ*, and *opy2Δ* mutants had a similar reduction in fMAPK pathway activity in glucose (**Fig 3A**); however, the *opy2Δ* mutant showed a stronger reduction than the *msb2Δ* or *sho1Δ* mutants in other environments [**Fig 3A,** p-values by Student’s t-test, galactose with pectin, p-value < 0.0002 to *msb2Δ* and < 0.00001 to *sho1Δ*, galactose with ethanol, p-value < 0.003 to *msb2Δ* and < 0.008 to *sho1Δ*]. The observation that Opy2p showed a stronger role than Msb2p or Sho1p is in agreement with previous results (*129*), tying Opy2p to Ste11p recruitment and activation (*52, 69, 142, 143*). We also noticed that in response to galactose with pectin, the *msb2Δ* mutant showed a larger reduction in fMAPK pathway activity than the *sho1Δ* mutant (**Fig 3A**, galactose with pectin, p-value < 0.00001 by Student’s t-test), whereas the *msb2Δ* and *sho1Δ* mutants showed similar reduction in galactose with ethanol Collectively, these results indicate that Msb2p, Sho1p and Opy2p can show similar or different requirements depending on the environment (**Fig 3B**, compare arrow sizes). Overall, these findings suggest that the stimuli tested may be sensed in different ways by different proteins.

We next found that some proteins were required in all the conditions tested as their deletion showed minimal reporter activity (**Fig 3A**, Ste50p, Ste11p, and Ste12p); whereas other relay proteins were partially or completely dispensable for signaling in certain contexts. For example, Msb2p, Sho1p, and Opy2p were partly dispensable in most environments, especially under maximally-inducing conditions (**Fig 3**, **A** and **B**, galactose with ethanol). In galactose with ethanol, Ste20p was also partially dispensable (**Fig 3**, **A** and **B**) and the Bem4p and Dig1p proteins were fully dispensable (**Fig 3**, **A** and **B**). We also surprisingly found that the two main transcription factors for the fMAPK pathway, Tec1p and Ste12p, which normally have the same phenotype, showed different requirements for fMAPK pathway activity. Ste12p was absolutely required for signaling in galactose with ethanol, but Tec1p was not (**Fig. 3**, **A** and **B**, p-value < 0.0004 by Student’s t-test comparing *tec1Δ* and *ste12Δ*). Collectively, these data show that the requirement for a subset of fMAPK pathway components changes depending on the condition tested.

### Components of the fMAPK pathway show variation in regulating filamentous growth

The fMAPK pathway regulates differentiation to the filamentous cell type (*28, 29*). Cell differentiation includes an elongation of cell shape, due to a delay in the G1 and G2 phases of the cell cycle (*41, 144, 145*), and a switch from axial budding (growth towards the mother cell) to distal budding [or growth away from the mother cell (*28, 54, 146, 147*)]. These features were observed in response to combinations of inducers by light microscopy and were represented as a ratio of cells undergoing filamentous growth compared to the total number of cells (% filamentous). A cell was considered filamentous if it exhibited an elongated cell morphology or distal budding pattern (see Material and Methods).

For wild-type cells, the percentage of filamentous cells (like for fMAPK pathway activity) increased from growth in glucose to growth in galactose media and from galactose to galactose with ethanol media (**Fig 4A**, representative images, **Fig 4B**, quantitation). We also noticed that the length of individual cells did not increase from galactose to galactose with ethanol (**Fig 4A**, WT, black arrows, compare GAL to GAL+E) even though fMAPK pathway activity increased between these environments (**Fig 3A**). These data suggest that the increase in fMAPK pathway activity may result from a higher number of cells in the population being stimulated, rather than a higher level of fMAPK pathway activity in individual cells. Moreover, the *dig1Δ* mutant was similar to wild-type cells in the maximally inducing condition (galactose with ethanol) for both fMAPK pathway activity (**Fig. 3A**) and the % filamentous cells (**Fig 4B**). Notably, the *dig1Δ* mutant did show more aberrant cell morphologies than wild-type cells (**Fig 4A**, compare red arrows), which may reflect that cells can tolerate fMAPK pathway activation by a natural stimulus compared to hyperactivation resulting from a genetic perturbation. This is possibly because Dig1p may have pleiotropic effects on cell morphology unrelated to its role in regulating fMAPK pathway activity.

**Figure 4.**
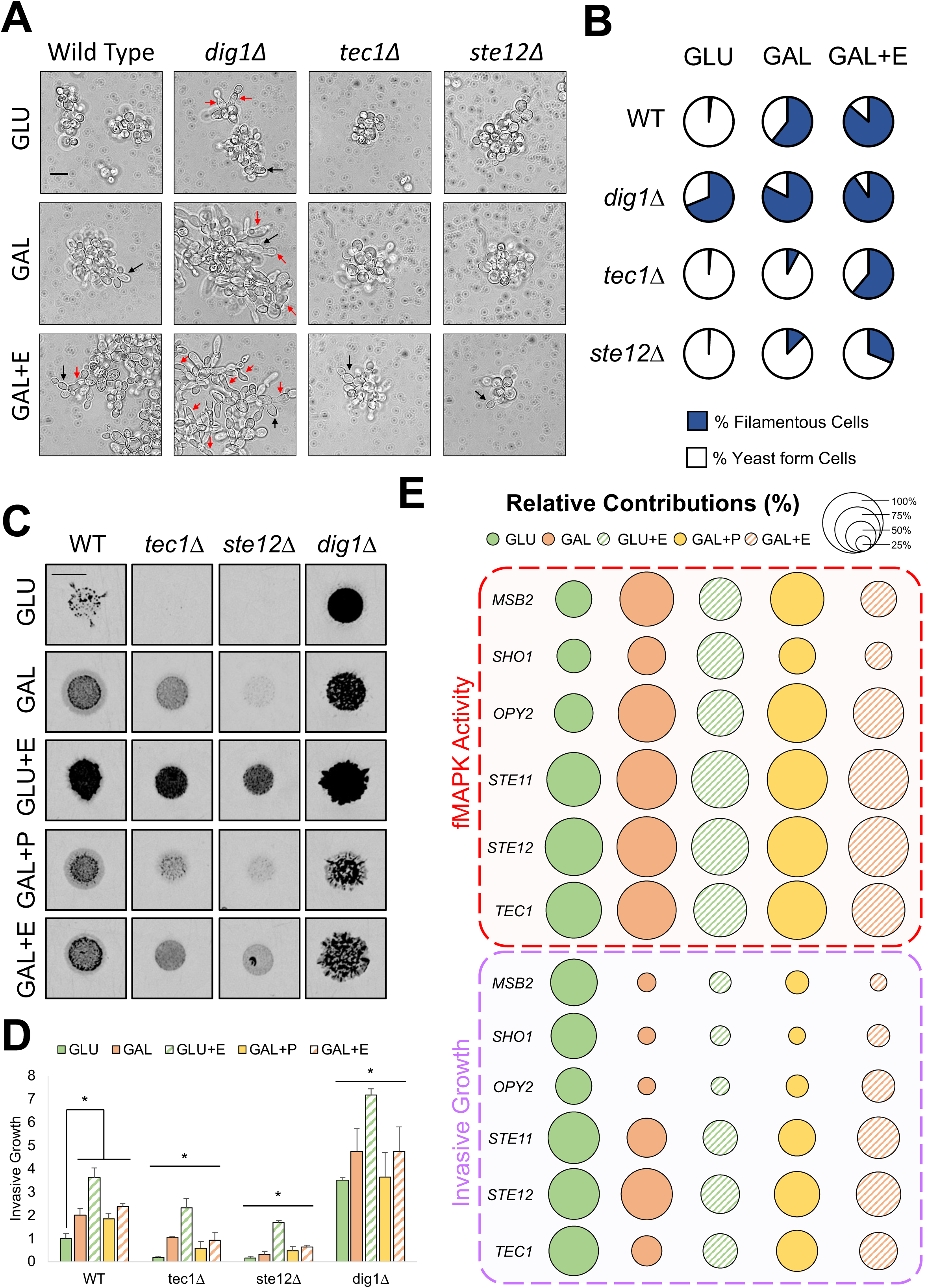
Environment-dependent roles for pathway components in filamentous growth. **A)** Wild-type (PC313) cells and the *ste12Δ* (PC5651), *tec1Δ* (PC7675) and *dig1Δ* (PC7676) mutants were examined by microscopy in indicated media and imaged. Microscopy images taken at 100X in indicated media. Bar, 10 um. Black arrow, filamentous cell. Red arrow, cell displaying aberrant morphology. **B)** Quantified % filamentous cells. Representative images in panel A. % filamentous cells (blue). % yeast-form cells (white). **C)** Plate-washing assay. Left, images, wild-type cells (PC313) and the *ste12Δ* (PC5651), *tec1Δ* (PC7675) and *dig1Δ* (PC7676) mutants were spotted for 5 d on indicated medium. Inverted images of invasive scars are shown. Bar, 0.5 cm. Before wash images of colonies are shown in *Fig S5A.* Results slightly differ from those published in (*141*) because they were performed for 5 d here instead of 7 d in the previous study. **D)** Quantitation of invasive growth in panel C. Average relative invasion across at least 3 replicates is reported, with wild-type values in GLU set to 1. Error bars represent standard deviation. Asterisk for wild-type values, p-value < 0.05 by Student’s t-test compared to indicated condition. Asterisk for mutants’ values, p-value < 0.05 by Student’s t-test compared to wild-type values from same condition. **E)** Relative % contribution for fMAPK pathway activity (top) or invasive growth (bottom) in the indicated conditions, where circle size represents the level of contribution. Larger circle, larger contribution. Contribution is determined as reported (*141*) based on the reduction in invasive growth relative to wild type converted to a percentage. Based on quantitation in *Fig S5C*.

For Ste12p and Tec1p in galactose with ethanol, the % filamentous cells (**Fig 4B**) matched fMAPK pathway activity (**Fig. 3A**), in that the *ste12Δ* mutant showed a stronger reduction in the % filamentous cells than the *tec1Δ* mutant. Surprisingly, in this context the *tec1Δ* and *ste12Δ* mutants showed some cell elongation and distal budding (**Fig 4A**, GAL+E, black arrows), which may come from another pathway that regulates filamentous growth. Thus, we identified a condition where Tec1p can be partly by-passed for both fMAPK pathway activity (**Fig. 3**) and filamentous growth (**Fig. 4**, **A** and **B**).

Filamentous growth can also be visualized by the plate-washing assay, where spotted cells washed with a stream of water show invasive growth (*38*). Invasive growth is normally tested on YPD [Yeast extract, peptone, dextrose] media, but here synthetic medium was used to match conditions where the *FRE* reporter was evaluated. The plate-washing assay showed that components of the fMAPK pathway were required for invasive growth more under some conditions than others (*Fig S5A* and *S5B*). For example, the plate-washing assay supported the finding that Ste12p plays a more critical role than Tec1p under several conditions (**Fig 4C**, e.g. GAL, quantitation in **Fig 4D**), which supports fMAPK pathway activity (**Fig 3A**) and % filamentous cells (**Fig 4B**) data above.

Invasive growth matched or showed differences compared to fMAPK pathway activity depending on the environment (direct comparison between fMAPK activity and invasion in *Fig S5, B-D*). For example, invasive growth (**Fig 4C** and **4D**, WT, compare down columns) matched fMAPK pathway activity (**Fig 3A**) based on the fact that galactose stimulated both phenotypes relative to glucose, and glucose with ethanol stimulated both phenotypes relative to glucose alone. Invasive growth differed from fMAPK pathway activity based on the fact that glucose with ethanol induced the most invasion (**Fig 4C** and **4D**, WT), whereas galactose with ethanol induced the most fMAPK pathway activity (**Fig 3A**, WT). In addition, combinations of galactose plus pectin or ethanol did not cause an increase in invasive growth beyond galactose alone (**Fig 4C** and **4D**, WT) as it did for fMAPK pathway activity (**Fig 3A**, WT). These results suggest that fMAPK pathway activity and invasive growth are separable phenotypes. This idea was not noted in our previous study (*141*), most likely because environments were not examined that maximally activate the fMAPK pathway. This result may suggest that fMAPK pathway activity and invasive growth become uncoupled at higher levels of pathway activation.

In support of the disconnect between fMAPK pathway activity and invasive growth, some fMAPK pathway components made strikingly different contributions toward those two phenotypes depending on the environment (**Fig 4E**, plate-washing assay images and quantification in *Fig S5*). For example, Msb2p, Sho1p, and Opy2p generally showed similar contributions to both the regulation of fMAPK pathway activity and invasion in glucose [**Fig 4E**, GLU, green, circle sizes are similar between fMAPK pathway activity (top) and invasive growth (bottom)] but showed striking differences in their contributions between the two phenotypes in inducing environments [**Fig 4E**, e.g. GAL, orange, circle sizes are much larger for fMAPK activity (top) than invasive growth (bottom)]. The downstream components (Ste11p, Ste12p, and Tec1p) were also variable in their contribution between the two phenotypes, but to a lesser degree (**Fig 4E**). Invasive growth may be less affected by fMAPK pathway perturbation than fMAPK pathway activity in some contexts because invasive growth can be independently regulated by other signaling pathways from the filamentous regulatory network (*27, 148*). Overall, the findings in this section suggest that environmental inducers stimulate invasive growth, cell differentiation, and fMAPK pathway activity, but not necessarily in the same manner, because fMAPK pathway components show variation between each other in some contexts.

### A key role for the Ras2-PKA pathway in the conditional regulation of the fMAPK pathway

Other pathways regulate filamentous growth, and a subset of the pathways also regulate the fMAPK pathway (*141, 149, 150*). Key mutants known to ablate the **RAS** (or Ras2p), **RIM**, **RTG**, **UPR**, **LIPID**, **HOG**, and **MIG-Glucose Repression (GR)** pathways were examined for fMAPK pathway activity under the conditions examined in this study. Most pathways played distinct roles in regulating the fMAPK pathway depending on the condition tested. For example, **RAS** was required under five conditions, whereas the **UPR** was only required under four conditions, and **RIM** was only required under 3 conditions (**Fig 5, A** and **B**). Moreover, some pathways played a positive role in regulating the fMAPK pathway in one environment and a negative role in another [**Fig 5, A** and **B**, e.g. **RTG**: positive, GLU, negative, GAL; **HOG**: positive, GAL+E, negative, GLU]. A ‘role reversal’ has previously been reported for **RTG** in multiple environments for invasive growth (*141*). Strikingly, **RAS** played a central role in regulating fMAPK pathway activity in galactose with ethanol, even more than Msb2p, Sho1p, Bem4p, or Ste20p (**Fig. 5A**). **RAS** also played a major role in invasive growth and showed a stronger contribution in response to ethanol (but not the absence of ethanol) than any component of the fMAPK pathway tested (*Fig S5*, *ras2Δ*). This result suggests that **RAS** is required for fMAPK pathway activity in this context, rather than functioning as an ancillary pathway that augments core pathway activity.

**Figure 5.**
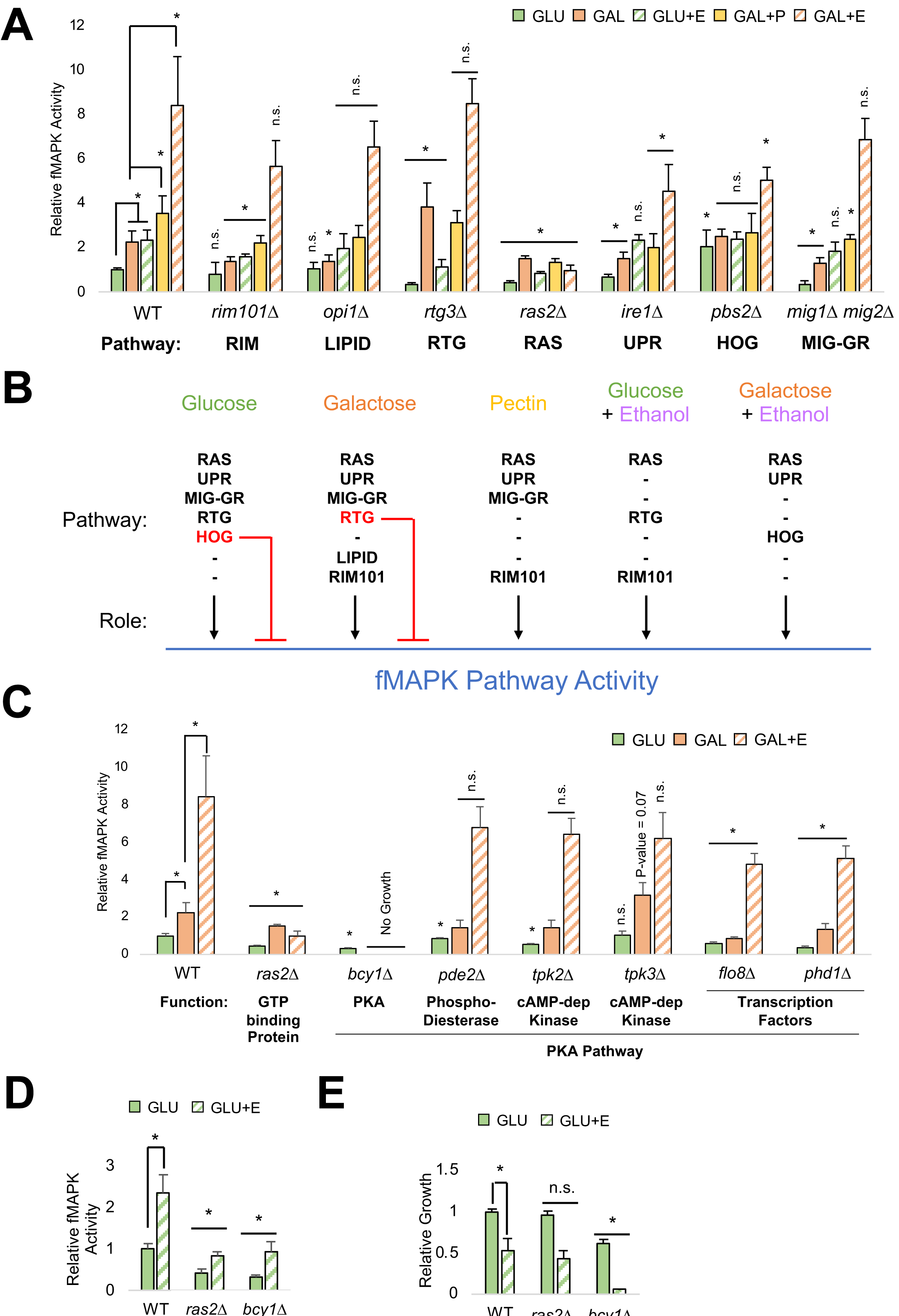
Role for the Ras2-PKA pathway in the conditional regulation of the fMAPK pathway. **A)** ß-galactosidase (*lacZ*) assays of mutants lacking pathways that regulate filamentous growth. RIM101 pathway [**RIM**] (*187, 188*); Phospholipid-biosynthesis-regulator-Opi1p [**LIPID**] (*141, 189*); Ras2p [**RAS**], cAMP-dependent protein kinase A (PKA) pathway (*28, 141, 149, 151, 153, 154*), Unfolded Protein Response [**UPR**, (*59*)], and AMPK-Mig1/2p-Glucose Repression pathway [**MIG GR**, (*29, 119*)]; the retrograde pathway [**RTG**] (*141, 190, 191*); and the High-Osmolarity-Glycerol pathway [**HOG**] (*89, 166, 192*). Wild-type (PC313) cells and the *rim101Δ* (PC7673), *opi1Δ* (PC7674), *rtg3Δ* (PC7677), *ras2Δ* (PC6222), *ire1Δ* (PC6048), *pbs2Δ* (PC5035), and *mig1Δ mig2Δ* (PC5187) mutants were grown in 2 mL synthetic media with indicated carbon source after 17 h. +P = + 1% pectin. +E = +3.85% ethanol. Average relative fMAPK pathway activity across at least 3 replicates is reported, with wild-type values in GLU set to 1. Error bars represent standard deviation. Asterisk for wild-type values, p-value < 0.05 by Student’s t-test compared to indicated condition. Asterisk for mutants’ values, p-value < 0.05 by Student’s t-test compared to wild-type values from same condition. Wild-type values are the same as shown in **Fig 3A**. **B)** Model of network-level regulation of fMAPK pathway activity. Black arrow, positive role. Red arrow, inhibitory role. Dash line, pathway does not serve a role in indicated condition.**C)** ß-galactosidase (*lacZ*) assays of the PKA pathway, which includes cAMP-dependent kinases Tpk2p and Tpk3p (*151, 193, 194*); transcription factors Flo8p and Phd1p (*53, 93, 195-197*); a phosphodiesterase Pde2p (*198–201*); a regulatory subunit of PKA, Bcy1p (*195, 202*). Performed same as panel A except wild-type cells (PC313) and the *ras2Δ* (PC6222), *bcy1Δ* (PC7870), *pde2Δ* (PC7872), *tpk2Δ* (PC7874), *tpk3Δ* (PC7869), *flo8Δ* (PC7865), and *phd1Δ* (PC7873) mutants were used. The *ras2Δ* mutant values are the same as shown in panel A. **D)** ß-galactosidase (*lacZ*) assays. Same as panel A except wild-type (PC313) cells and the *ras2Δ* (PC6222) and *bcy1Δ* (PC7870) mutants were used. The *ras2Δ* mutant values are the same as shown in panel A. The *bcy1Δ* mutant value in GLU is the same as shown in panel C. **E)** Relative growth of wild-type (PC313) cells and the *ras2Δ* (PC6222) and *bcy1Δ* (PC7870) mutants in 2 mL synthetic media with indicated carbon source after 17 h. Average relative growth across at least 3 replicates is reported, with wild-type GLU values set to 1. Error bars represent standard deviation. Asterisk for wild-type values, p-value < 0.05 by Student’s t-test compared to indicated condition. Asterisk for mutants’ values, p-value < 0.05 by Student’s t-test compared to wild-type values from same condition.

**RAS** may regulate the fMAPK pathway by direct interactions between fMAPK pathway components and the GTPase Ras2p (*151, 152*) or through Ras2p regulation of the protein kinase A (PKA) pathway (*153, 154*). Strains lacking PKA components were generated and tested for fMAPK pathway activity in response to galactose with ethanol (**Fig 5C**). Some PKA mutants showed a reduction (*tpk2Δ* and *pde2Δ*, GLU) or increase (*tpk3Δ*, GAL) in fMAPK pathway activity in one environment (**Fig 5C**). Others showed a decrease across all environments tested (*flo8Δ, phd1Δ,* **Fig 5C**). The deletion of the gene encoding Bcy1p showed a strong and comparable reduction as the *ras2Δ* mutant for fMAPK pathway activity in glucose (**Fig 5D**, green), though it also showed reduced growth (**Fig 5E**, green). The *bcy1Δ* mutant did not grow in galactose, so we tested the mutant in glucose with ethanol, where it showed a similar result (**Fig 5, D** and **E**). Thus, both Ras2p and PKA pathway disruptions reduced fMAPK pathway activity in response to ethanol. Two-hybrid tests did not identify an interaction between Ras2p and fMAPK pathway components (*Fig S6*); however, Ras2p may directly regulate the fMAPK pathway as it interacts with Cdc24p (*155*) and the polarity adaptor Bem1p (*77–79*) which both regulate the fMAPK pathway (*77, 80*). Thus, the Ras2-PKA pathway is important to fMAPK pathway activity in response to ethanol including conditions (e.g. galactose with ethanol) where the pathway functions near-maximal levels.

## DISCUSSION

*S. cerevisiae*, like the pathogens *C. albicans* and *U. maydis*, undergo a type of filamentous growth response that is regulated by a MAPK pathway when exposed to specific environments. Unlike pathogens which are commonly studied in their hosts, MAPK pathway function in *S. cerevisiae* has been studied mostly under standard laboratory conditions. To examine the fMAPK pathway from an ecological perspective, we tested the effect of compounds that may be encountered by *S. cerevisiae* in the wild (e.g. plant-derived compounds) on fMAPK pathway signaling and filamentous growth. Three inducers of the fMAPK pathway were identified: pectin, galactose, and ethanol. Moreover, combinations of inducers (galactose with ethanol) dramatically activated the pathway to near-maximal levels. These diverse signals transduce through the fMAPK pathway through different subsets of pathway components and surprisingly revealed a difference between the transcription factors Ste12p and Tec1p. Maximal signaling also required the Ras2-PKA pathway. Thus, an ecological perspective led to new insights about signaling pathway induction and regulation.

The fact that pectin can induce the fMAPK pathway suggests a closer connection between yeast and plants than previously appreciated. One possibility is that this connection represents a role for the pathway in plant-environment recognition. Because the fMAPK pathway controls expression of the cell’s major pectinase (*41*), detecting pectin may lead to the break down of plant material to release sugars to scavenge. Sugars released from plant tissues can consist of preferred carbon sources (e.g. glucose and fructose) and non-preferred carbon sources (e.g. galactose, sucrose, and maltose). Glucose is preferentially used and prevents the use of non-preferred carbon sources in yeast (*119, 156, 157*). We found that glucose also prevents pectin induction of the fMAPK pathway. This may be because filamentous growth evolved to occur only in the absence of a preferred carbon source. Additionally, we found that galacturonic acid, a compound that can be introduced into the environment when other microbes forage and break down pectin (*123–126*), inhibited growth and fMAPK pathway activity. This may suggest galacturonic acid is involved in microbial competition by *S. cerevisiae* competitors.

We also found that galactose is more effective at activating the fMAPK pathway than glucose limitation or other non-preferred carbon sources. The differences between effects on fMAPK pathway activity by glucose, maltose, galactose, and sucrose may be because the sugars are metabolized differently, require different enzymes and permeases, and enter glycolysis in different ways (*158*). Because galactose is found in abundance in natural habitats [e.g. fruits and forests], this sugar may signal that cells have encountered a nutrient-replete environment, such as fruit, from which nutrients can be obtained. Interestingly, galactose and pectin induce the secretion of pectolytic enzymes in distantly related filamentous fungi, like *Neurospora crassa*, where glucose also inhibits pectin induction (*159–162*).

Carbon sources from the plant are used in glycolysis to produce energy, which in *S. cerevisiae* will produce ethanol as a byproduct; therefore, during the utilization of nutrients, ethanol accumulates and can indicate the decline in available nutrients. Additionally, ethanol acts as a quorum-sensing molecule (*35*), which are indicators of cell density across microbes (*36, 163*). We show here that ethanol, a known inducer of filamentous growth (*33*), also stimulates the fMAPK pathway. The fact that ethanol can induce fMAPK pathway activity in glucose suggests it represents an independent signal from nutrient levels. It may instead be a signal to cells that scavenging will be required for a growing population. In addition, ethanol induced the fMAPK pathway to higher levels in galactose, which suggests it may also act to amplify the low-nutrient signal. Thus, ethanol may integrate two important signals, low nutrients and high cell density.

It is not yet clear how pectin, galactose, and ethanol are “sensed” by the fMAPK pathway. Pectin may be sensed by associating with Msb2p at the cell surface, since intriguingly some mammalian mucins expressed in gut tissue can interact with pectin to facilitate the digestive process (*164, 165*). Another possibility is that large pectin molecules may act as a scaffold by interacting with several membrane-bound proteins (e.g. Msb2p, Sho1p, and Opy2p) and bring signaling complexes together. How pectin activates Msb2p and Sho1p is of interest because Msb2p-like and Sho1p-like proteins head MAPK pathways that are involved in host recognition and invasion in plant pathogens like *U. maydis* (*24, 25*), *Fusarium oxysporum* (*22*), *Magnaportha oryzae* (*23*), and *Colletotrichum gloeosporioides* (*21*).

Galactose signaling requires galactose metabolism. This may suggest that one of the components of the galactose utilization pathway may interact with the cytosolic domains of Msb2p and/or Sho1p. Alternatively, galactose metabolism may feed into the pathway because the glycosylation of Msb2p becomes reduced, which leads to elevated processing of Msb2p and fMAPK pathway induction (*59, 166*). Ethanol may be sensed through its ability to denature proteins or disrupt the structure of cell membranes (*167, 168*), which is where the fMAPK pathway sensors are located. Understanding how these different signals are sensed is of future interest because findings may apply generally to many fungal species that contain similar sensory pathways.

We previously measured the activity of the fMAPK pathway under several conditions that included variations in carbon, nitrogen, and phosphate levels and sources (*141*). Most conditions caused a modest induction of fMAPK pathway activity, and no condition induced the pathway to greater than half-maximal values. We proposed that the fMAPK pathway may never reach maximum levels by normal processes to protect the cells from detrimental phenotypes that occur in hyperactive MAPK states, such as defects in cell morphology and budding (*58, 150, 169*). Surprisingly, here, the ecologically-relevant stimuli in combination were able to maximally activate the fMAPK pathway. This did not cause wild-type cells to be ‘sick’ like in hyperactive mutants (e.g. *dig1Δ*). These findings imply that maximal activation of the fMAPK pathway can occur in the wild and may suggest a beneficial state for the cells in some contexts. Maximal activation appears to occur by inducing the pathway across more cells in the population, rather than stronger activation in individual cells. This maximally-activated state also revealed that fMAPK pathway activity correlates to some phenotypes (e.g. cell differentiation) but is separable from others (e.g. invasive growth) in some contexts. Broadly speaking, these findings could suggest two things relative to other MAPK pathways and organisms. First, this could imply that MAPK pathways activity and output phenotypes do not directly correlate as one might predict at different levels of activation. Second, the prior techniques of studying MAPK states through the introduction of hyperactive alleles of pathway components or the deletion of negative regulators may not fully represent what cells do when a MAPK pathway is activated. Therefore, the ability to activate the fMAPK pathway to high levels with commonly encountered stimuli without generating obvious morphological abnormalities represents a new and powerful tool for molecular dissection of pathway function.

This ability to naturally activate the fMAPK pathway to high levels allowed us to identify conditional roles for fMAPK pathway components. These include our findings that, in galactose with ethanol, Dig1p does not inhibit the fMAPK pathway, which suggests that under this condition the pathway is maximally activated. Under this condition, we also found that the scaffold Bem4p was dispensable, the sensors Msb2p, Sho1p, and Opy2 were partly dispensable, and the major PAK for the pathway, Ste20p, was partly dispensable. Ste20p has previously been found to be dispensable for ethanol-dependent filamentous growth compared to other pathway components (*33*), can be bypassed by activated versions of Cdc42p (*170*), and is partly redundant in a related signaling pathway (*65*). The Ras2-PKA pathway was also found to play a strong role in regulating fMAPK pathway activity in the maximally inducing environment, even more so than canonical sensory components (Msb2p, Sho1p, Ste20p, and Bem4p). Perhaps Ras2p bypasses these sensory proteins in a condition-specific manner.

Although there is some evidence of Tec1p regulatory effects on filamentous growth independent of Ste12p (*92, 171, 172*), the two transcription factors are thought to work mostly in concert during filamentous growth to induce target genes due to the formation of a heterodimer to regulate gene expression (*173*), and the fact that they play equal roles in many environments. Surprisingly, we found a role for Ste12p in the regulation of filamentous growth that can partly bypass Tec1p. This might suggest that Ste12p regulates target genes that are relevant to filamentous growth independently of Tec1p. The different environments may lead to separate regulatory mechanisms of the transcription factors by other proteins. For example, Ste12p associates with another transcription factor that is an essential gene, Mcm1p (*173*), and Ste12p is phosphorylated by Cdk8p in some contexts which reduces protein stability (*174*). Tec1 can also be independently regulated by SUMOylation which effects its regulation in invasive growth (*175*).

In conclusion, studying *S. cerevisiae* from an ecological perspective has led to new insights about signaling pathway induction and regulation. These findings may apply to human pathogens, like *C. albicans* where fMAPK pathway components are evolutionarily conserved (*176–178*), and suggest a focus on drug targeting core components (e.g. CaSte50, CaSte11 and the Ste12 homolog, Cph1). These findings may also apply more generally to many systems, including other animal and plant pathogens who interact with hosts in unique environments through MAPK signaling pathways. Broadly speaking, studying signaling pathways in diverse and meaningful contexts may reveal environment-dependent roles for pathway components that may broaden our understanding of health, disease, and evolution.

## MATERIALS AND METHODS

### Yeast Strains and Plasmids

Yeast strains are listed in *Table S1*. The p*FRE-lacZ* plasmid is used to measure the transcriptional activity of the fMAPK pathway (*118*) and was provided by H. Madhani (UCSF). Gene deletions were made in the ∑1278b strain background (*28*) through homologous recombination, constructed using an antibiotic resistance marker (neurotactin or gentamycin) amplified by polymerase chain reaction (PCR) and introduced into yeast by lithium acetate transformation as described (*179*). All gene deletions were verified by PCR amplification and gel electrophoresis of deletion site and by phenotype when possible. PCR primers for homologous recombination can be found in *Table S2*.

### Media

Synthetic complete medium (GLU, FRU, GAL, SUC, MAL, GLY): 0.67% yeast nitrogen base without amino acids, amino acids, and one of the following: 2% dextrose, 2% fructose, 2% galactose, 2% sucrose, 2% maltose, or 2% glycerol respectively (minus uracil when selecting for p*FRE-lacZ* plasmid). For some experiments with GLU medium, the dextrose concentration varies as indicated. <u>+P</u>: For media with pectin, a 1% solution was made using citrus pectin from Spectrum catalog #PE100 (https://www.spectrumchemical.com/pectin-citrus-usp-pe100); <u>YPD</u>: 1% yeast extract, 2% peptone, 2% dextrose; <u>YPGAL</u>: YPD with 2% galactose instead of dextrose. When solid media was made, 2% agar was used.

### Measurement of fMAPK pathway Activity

The fMAPK pathway activity was analyzed by the ß-galactosidase (*lacZ*) assay as previously described (*180, 181*) using an indicated transcriptional reporter [p*FRE*-*lacZ*] as the readout of fMAPK pathway activity. Cells were grown in indicated medium (without uracil when maintaining selection for plasmids). Cells were grown to timepoints indicated and harvested by centrifugation and stored at -80° for at least 30 minutes. Subtle variation in fMAPK pathway induction or growth by inducers can be noted across some figures. This is due to differences between experiments, as indicated in figure legends, related to different times cells were grown or different volumes of medium. The ß-galactosidase (lacZ) assay was performed with at least three biological replicates where the average is reported and error bars represent standard deviation. Differences in values from a previous study (*141*) for wild type and the *ras2Δ*, *rim101Δ*, and *opi1Δ* mutants may be due to differences in starting cell density and incubation times.

### Quantification of Phenotypes

Images of cells were taken at 100X magnification by microscopy using differential interference contrast (DIC) imaging with a Zeiss Axioplan 2 microscope. Digital images were acquired with the Axiocam MRm camera. Some photos were taken by an iPhone 13 (Apple) through the microscope lens. For image analysis, Axiovision 4.4 software was used.

For cell differentiation quantification, cells were observed at 100X. At least 85 cells were examined for each strain. Cells that showed elongation and/or cells that budded distally were considered filamentous, whereas cells that were round and budding axially (back toward mother cell) were considered yeast-form. The filamentous cells and yeast-form cells were expressed as a percentage of total cells.

To measure invasive growth, cells were spotted onto indicated medium and grown for 5 d. The plate-washing assay was performed as described (*54, 182*). Invasive growth was quantified as described using Image Lab https://www.bio-rad.com/en-us/product/image-lab-software?IKRE6P5E8Z to calculate volume/area x 10000 (*141*). Invasion values were averaged across at least three biological replicates and error was determined by standard deviation.

Pectin breakdown was determined by viscosity using a drop assay after cells were grown in 10 mL of indicated medium with 1% pectin for 20 h shaking at 30°. Viscosity was measured by dropping a screwcap Eppendorf tube filled with glass beads (weighing 4.4g) cap side down directly into the test tube, which was placed at a 45° angle. The time for the Eppendorf tube to reach the bottom was recorded for each strain. Viscosity was averaged across three biological replicates and error was determined by standard deviation. The breakdown of pectin in mandarin oranges was tested by growing the indicated strains for 24 h in 200 mL YPGAL liquid cultures shaking at 30°. Cells were spun down by centrifugation and cell supernatants were harvested. Mandarins were purchased commercially from Sunrays (https://sunraysfruits.com/products/mandarins/). The mandarin was sterilely peeled and wedges were separated. One mandarin wedge and one piece of peel was placed in 50 mL supernatants of indicated strains and left at 30° for 24 h. Wedges and peels were removed and placed under a gentle stream of water and washed. Images before and after treatment were captured using an iPhone 13 (Apple).

### Two-Hybrid Analysis

Two-Hybrid analysis (*183*) was performed to identify protein interactions as described in (*80, 184*). The assay was done by transforming two-hybrid constructs into the yeast strain background PJ69-4a using the pGAD-C1 – pGBDU-C1 system described in (*185*). Briefly, cells were spotted onto SD – Ura – Leu to maintain plasmid selection and to act as a control for growth. Cells were also spotted onto SD – Ura – Leu – His to assess the *LYS2::GAL1-HIS3* growth reporter. Positive control of a Cdc42p-Bem4p interaction is described in (*77, 80, 186*). Plasmid constructs used here are described in (*80*) for *RAS2*, *RAS2^G19V^*, *CDC24*, *STE11*, *STE50*, *KSS1*, *STE7*; in (*77*) for *BEM4, CDC42*, *BEM1*; in (*129*) for *OPY2*-tail; and in (*184*) for *STE20*.

## ABBREVIATIONS

AMPK: AMP-dependent protein kinase
fMAPK: filamentous growth mitogen-activated protein kinase pathway
FRU: fructose
GAL: galactose
GAP GTP: ase activating protein
GEF: guanine nucleotide exchange factor
GLU: glucose
GLY: glycerol
HOG: High-osmolarity-glycerol
IAA: indoleacetic acid
MAL: maltose
MAPK: mitogen-activated-protein kinase
MAPKK: mitogen-activated-protein kinase kinase
MAPKKK: mitogen-activated-protein kinase kinase, kinase
MIG GR: AMPK-Mig1p/2p-Glucose Repression; n.s., not significant
PAK: p21-activated kinase
PCR: polymerase chain reaction
PKA: Protein-kinase A
RTG: retrograde pathway
UPR: unfolded protein response
Y2H: yeast-2-hybrid
YPD: yeast extract, peptone, dextrose
YPGAL: yeast extract, peptone, galactose

## ACKNOWLEDGEMENTS

Thanks to Hiten Madhani for sharing the *pFRE-lacZ* reporter. Thanks to the past and present members of the Cullen Lab. Thanks to Kate and Avery Vandermeulen and Julie Aquino for their support.

## SUPPLEMENTAL FIGURE LEGENDS

**Figure S1.**
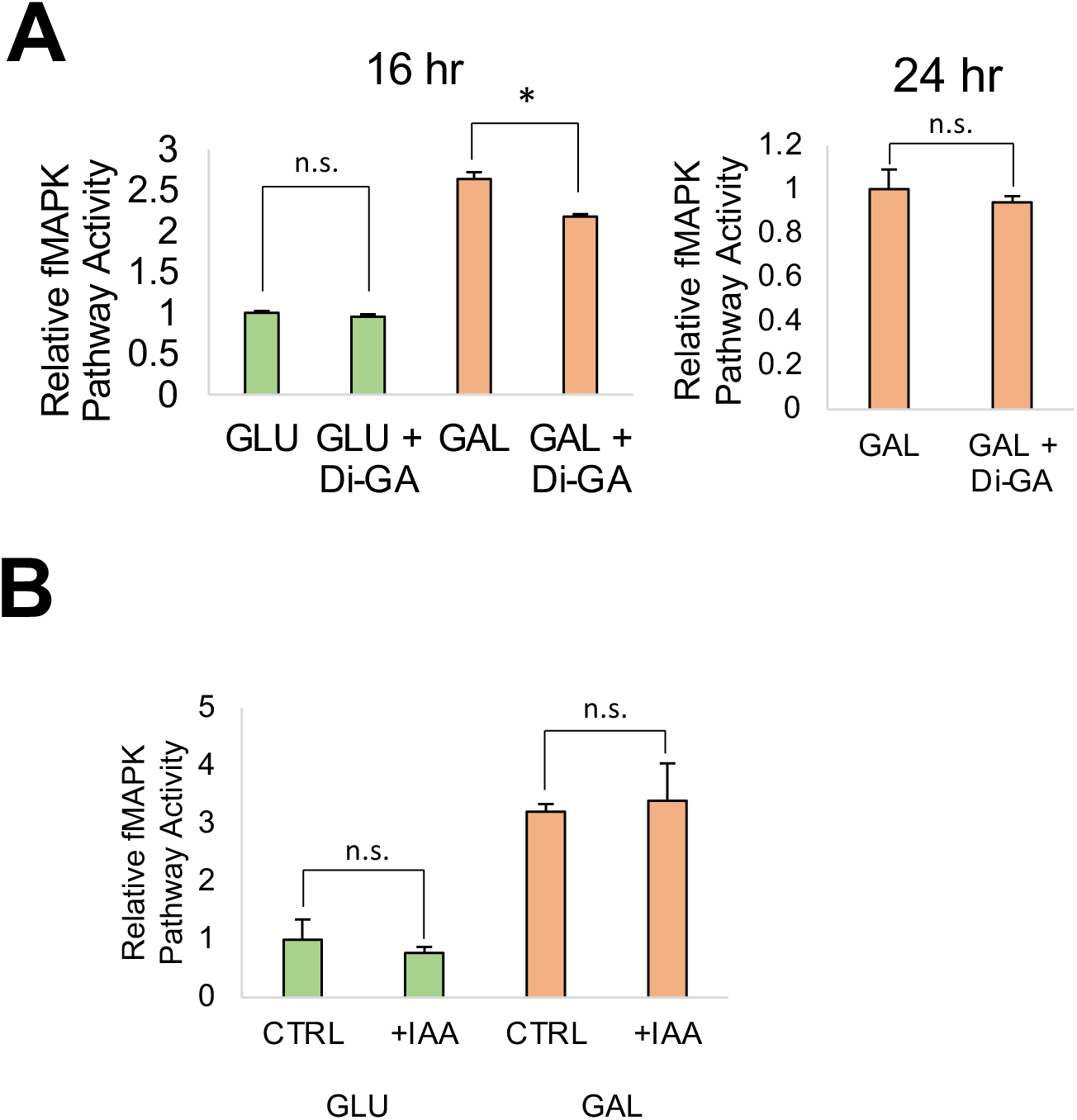
fMAPK pathway activity for galacturonic acid, di-galacturonic acid, and IAA. ß-galactosidase (*lacZ*) assays were performed in wild-type (PC313) cells. **A)** Cells were grown in 5 mL synthetic media with indicated carbon source after 16 h (left) or 24 h (right) and +3 mg of di-galacturonic acid as indicated. Average relative fMAPK pathway activity across at least 3 replicates is reported, with GLU values set to 1 (left) or GAL values set to 1 (right). Error bars represent standard deviation. Asterisk, p-value < 0.05 by Student’s t-test for the indicated comparisons. **B)** Cells were grown in 5 mL synthetic media with the indicated carbon source after 20 h and +0.1 mg of IAA dissolved in NaOH as indicated. For the control (CTRL), an equal amount of NaOH alone was added to the media. Average relative fMAPK pathway activity across at least 3 replicates is reported, with CTRL values in GLU set to 1. Error bars represent standard deviation. Asterisk, p-value < 0.05 by Student’s t-test for indicated comparisons.

**Figure S2.**
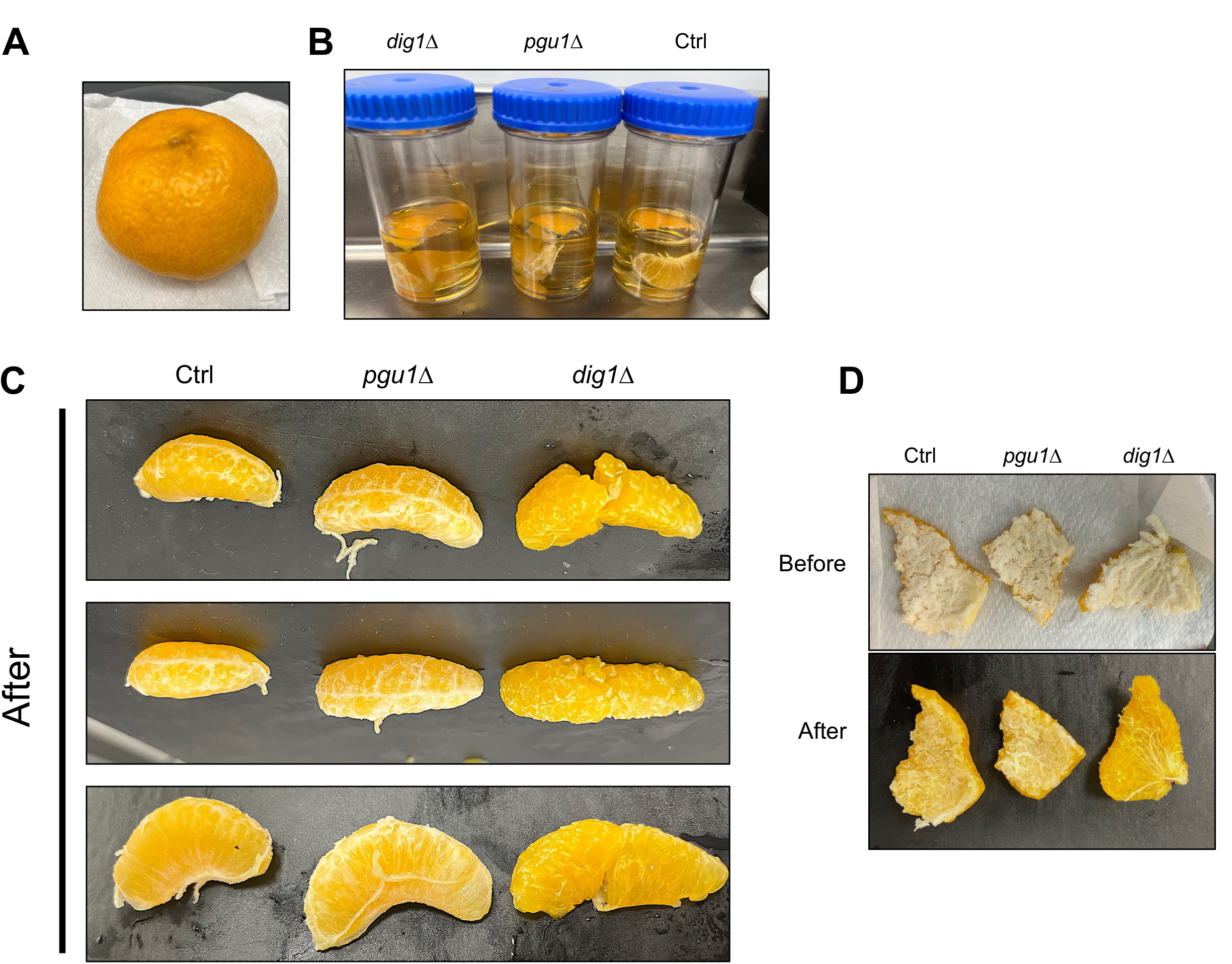
Examples of mandarin orange experiments. Mandarin orange wedges were submerged for 24 h in supernatants of *pgu1Δ* (PC7833) and *dig1Δ* (PC7676) mutants. Supernatants were derived from 24 h cultures in YPGAL. Ctrl, YPGAL media with no cells added. **A)** Image of mandarin orange. **B)** Image of mandarin wedges and peels submerged in cell supernatants at time zero. **C)** Images of wedges after submersion after 24 h. Three examples are shown. **D)** Images of peels before (top) and after (bottom) 24 h incubation with cultured supernatants.

**Figure S3.**
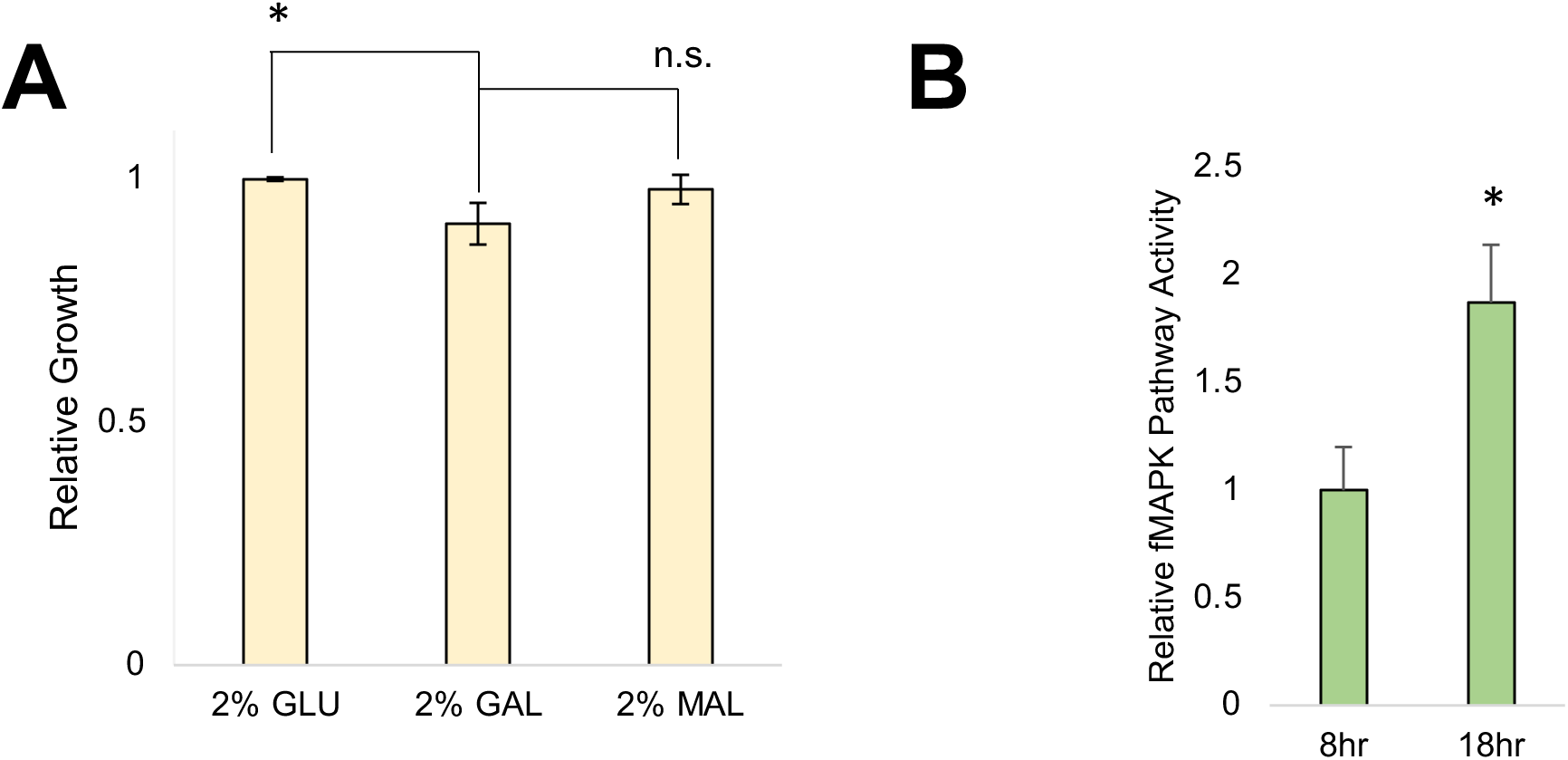
Glucose depletion stimulates the fMAPK pathway. **A)** Relative growth of wild-type cells (PC313) grown in 5 mL of synthetic media with indicated carbon source for 24 h. Average relative growth determined by OD_600_ across at least 3 replicates is reported, with GLU values set to 1. Error bars represent standard deviation. Asterisk, p-value < 0.05 by Student’s t-test compared to 2% GAL. **B)** Wild-type cells (PC313) were grown in 5 mL synthetic media with glucose for 8 h or 18 h. Average relative fMAPK pathway activity of at least 3 replicates are reported with 8 h GLU values set to 1. Error bars represent standard deviation. Asterisk, p-value < 0.05 by Student’s t-test compared to 8 h GLU values.

**Figure S4.**
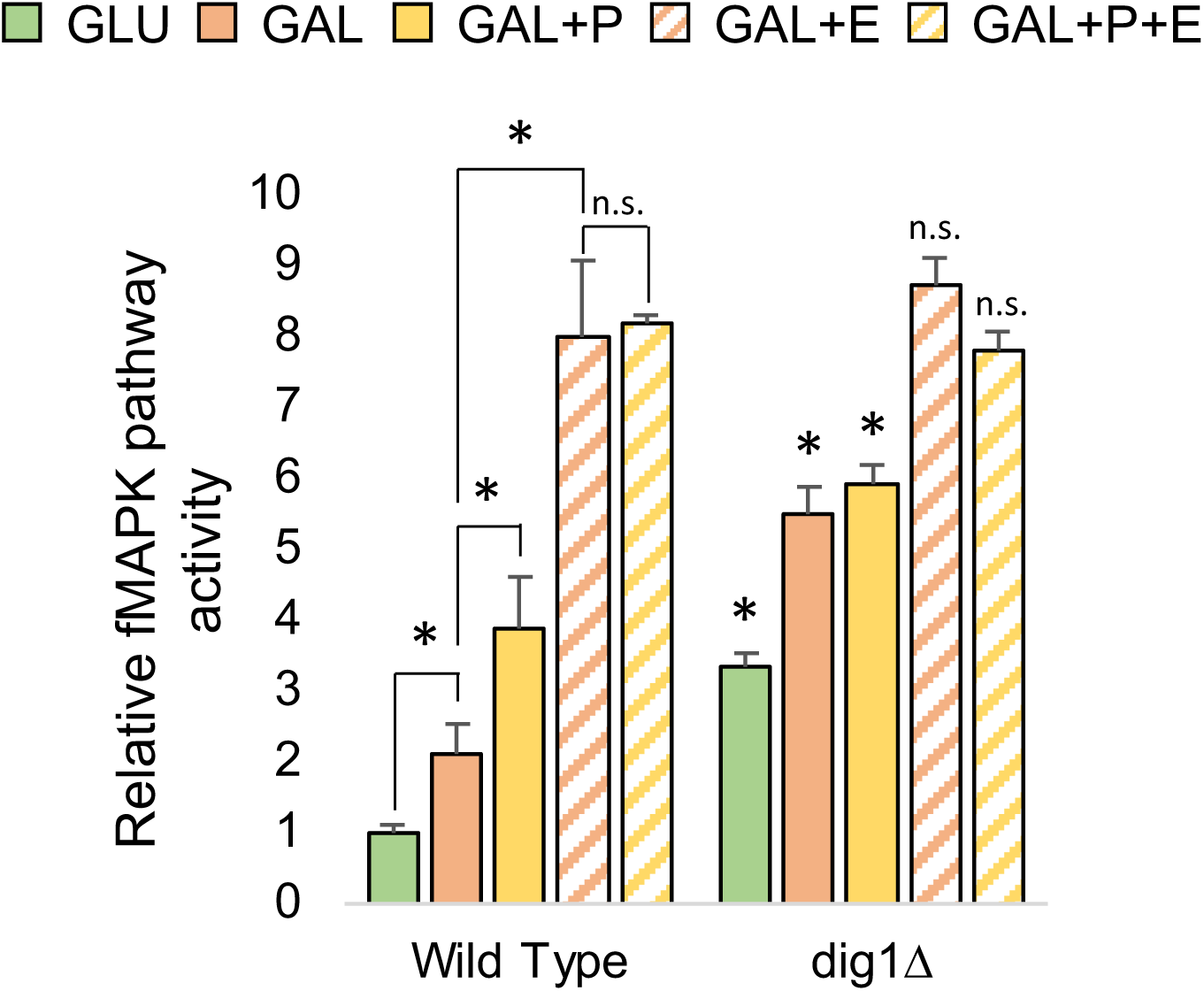
Combinatorial effects of inducers on fMAPK pathway activity. ß-galactosidase (lacZ) assays. Wild-type (PC313) cells and the *dig1Δ* (PC7676) mutant were grown in 2 mL synthetic media with indicated carbon source for 24 h. +P = + 1% pectin. +E = +3.85% ethanol. Pectin with ethanol was not tested because cells failed to grow due to a lack of carbon source, combined with the growth inhibition caused by ethanol. Average relative fMAPK pathway activity across at least 3 replicates is reported, with wild-type GLU values set to 1. Error bars represent standard deviation. Asterisk for wild-type values, p-value < 0.05 by Student’s t-test compared to indicated condition. Asterisk for *dig1Δ* values, p-value < 0.05 by Student’s t-test compared to wild-type values from same condition.

**Figure S5.**
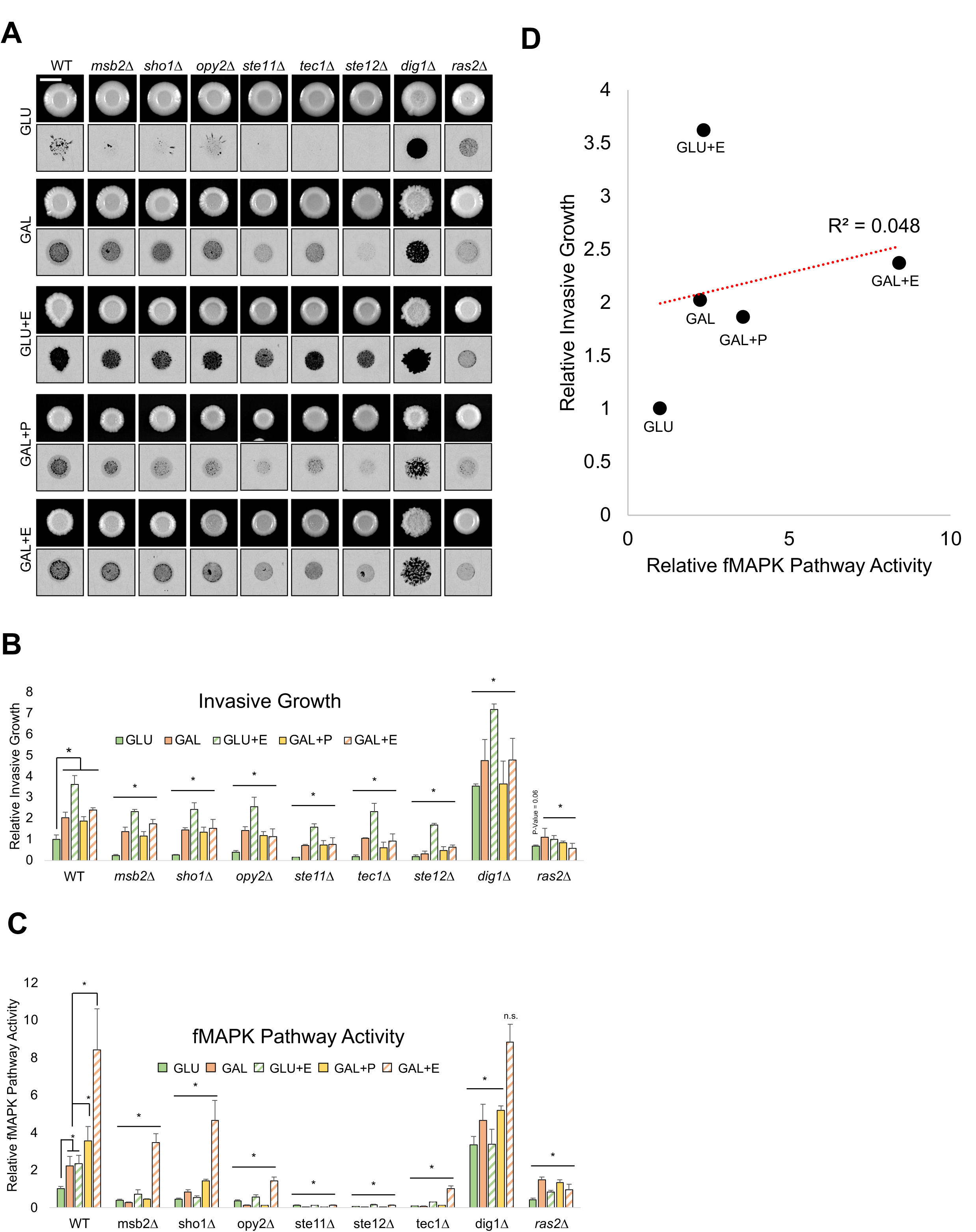
Full analysis of the plate-washing assay. **A)** Plate-washing assay. Wild-type (PC313) cells and the *msb2Δ* (PC961), *opy2Δ* (PC3894), *sho1Δ* (PC5026), *ste11Δ* (PC5024), *ste12Δ* (PC5651), *tec1Δ* (PC7675), *dig1Δ* (PC7676), and *ras2Δ* (PC6222) mutants were spotted for 5 d on the indicated medium. Top row, before wash. Bottom row, inverted images of invasive scars after wash. Bar, 0.5 cm. **B)** Quantitation of the data shown in panel S5A. Average relative invasion across at least 3 replicates is reported, with wild-type values in GLU set to 1. Error bars represent the standard deviation. Asterisk for wild-type values, p-value < 0.05 by Student’s t-test compared to indicated condition. Asterisk for mutants’ values, p-value < 0.05 by Student’s t-test compared to wild-type values from same condition. **C)** The same fMAPK pathway activity data from **Fig 3A** for comparison to invasive growth. **D)** fMAPK pathway activity compared to invasive growth for wild-type cells on indicated media reveals no correlation, R^2^ = 0.048. Data values are repeated from panels B and C.

**Figure S6.**
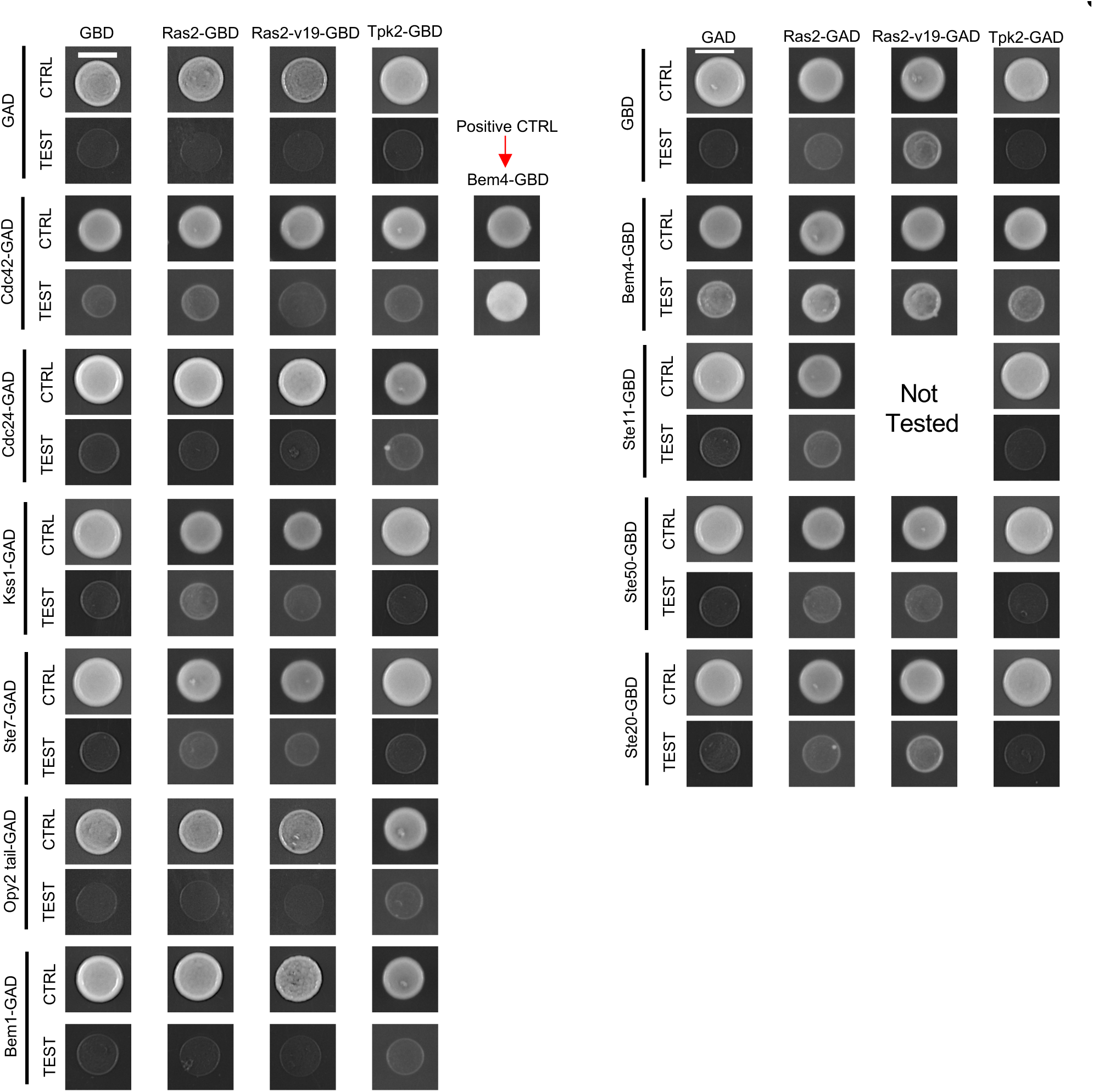
Two-hybrid analysis. Y2H analysis between Ras2p, a hyperactive version (Ras2^G12V^), and the PKA pathway kinase, Tpk2p, with components of the fMAPK pathway. Red arrow, interaction between Bem4p and Cdc42p (*80*) was used as a positive control. CTRL, SD-Ura-Leu media supplemented with HIS. TEST, SD-Ura-Leu media not supplemented with HIS. Growth on TEST plate represents an interaction.

**Table S1.**
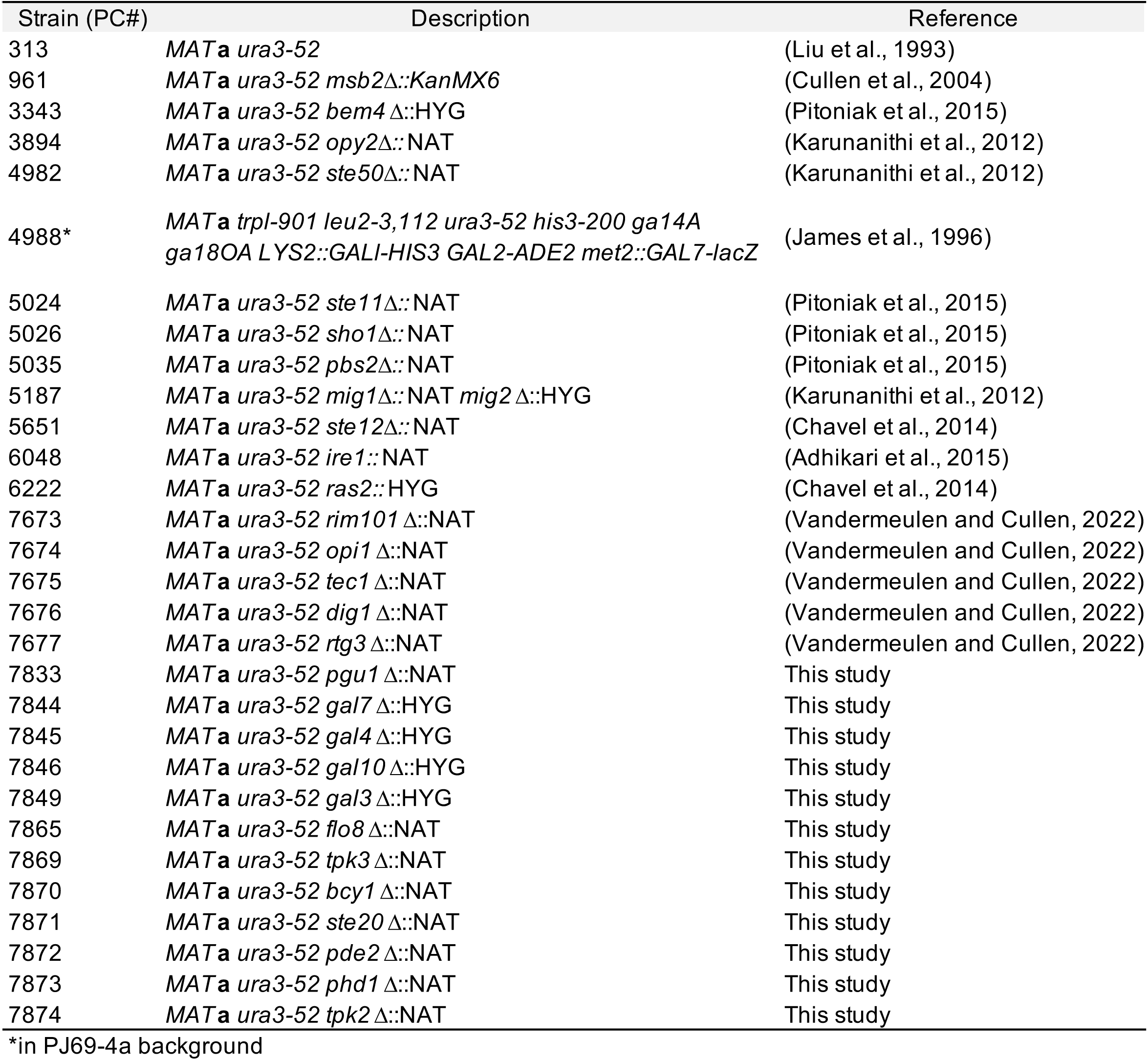
Yeast strains used in this study.

**Table S2.**
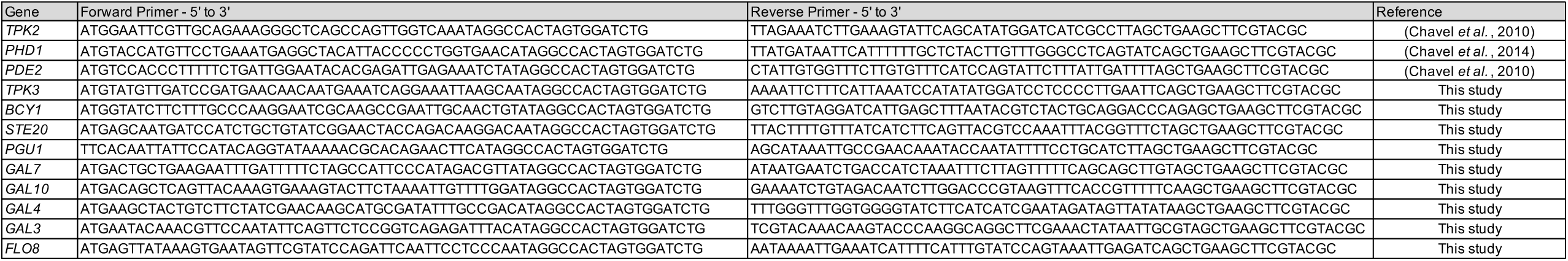
Primers used in this study to generate deletion mutants.

